# WRN Inhibition Leads to its Chromatin-Associated Degradation Via the PIAS4-RNF4-p97/VCP Axis

**DOI:** 10.1101/2023.12.08.570895

**Authors:** Fernando Rodríguez Pérez, Dean Natwick, Lauren Schiff, David McSwiggen, Melina Huey, Alec Heckert, Mandy Loo, Rafael Miranda, Huntly Morrison, Jose Ortega, Renee Butler, Kylie Cheng, John Filbin, Zhengjian Zhang, Eric Gonzalez, Rand Miller, Yangzhong Tang, Jaclyn Ho, Daniel Anderson, Charlene Bashore, Steve Basham

## Abstract

Synthetic lethality, the concept in which the co-occurrence of two genetic events leads to cell death while either single event alone does not, is an attractive strategy for targeted cancer therapies. A recent example of synthetic lethality as a therapeutic paradigm is the observation that cancer cells with high levels of microsatellite instability (MSI-H) are dependent on the Werner (WRN) RecQ helicase for survival. However, the mechanisms that regulate WRN spatiotemporal dynamics are not fully understood. In this study, we used our single molecule tracking (SMT) platform in combination with a recently disclosed WRN inhibitor to gain insights into WRN’s dynamic localization within the nuclei of live cancer cells. We observe that WRN inhibition results in the helicase becoming trapped on chromatin, requiring p97/VCP for extraction and shuttling to the proteasome for degradation. Interestingly, this sequence of events resulting in WRN degradation appears to be MSI-H dependent. Using a phenotypic screen, we identify the PIAS4-RNF4 axis as the pathway responsible for WRN degradation and show that co-inhibition of WRN and SUMOylation has an additive toxic effect in MSI-H cells. Taken together, our work elucidates a novel regulatory mechanism for WRN. Gaining a deeper understanding into this regulatory pathway for WRN can aid in the identification of new high value targets for targeted cancer therapies.

## Main

Werner Syndrome is a rare genetic condition caused by mutations in the *WRN* gene. It is marked by accelerated aging and a predisposition to a variety of cancers (Opresko et al., 2003). The *WRN* gene encodes a RecQ helicase that plays a critical role in genomic integrity and is unique amongst RecQ helicases as it possesses an exonuclease domain in addition to its helicase domain (Croteau et al., 2014). WRN typically resides in the nucleoli of cells but undergoes DNA damage-induced translocation to the nucleoplasm to perform its repair functions, resolving a diverse array of DNA substrates including D-loops, replication forks bubble structures, Holliday junctions and other secondary structures. (Constantinou et al., 2000; von Kobbe and Bohr 2002; Bendtsen et al., 2014). Additionally, WRN is required for telomere maintenance, a necessary process for maintaining stem cell cellular homeostasis (Shen and Loeb 2001).

Synthetic lethality has emerged as an appealing approach to target cancer cells while minimizing collateral damage to otherwise healthy cells and tissue (O’Neil et al., 2017). Synthetic lethality occurs when there is simultaneous disturbance of two essential biological events leading to cell death, which would not occur in the presence of one genetic disruption alone. To highlight the importance of WRN in DNA repair, WRN has been identified as a synthetic lethal target in microsatellite instability high (MSI-H) tumor types that are deficient in mismatch repair (MMR) pathways (Chan et al., 2019; Kategaya et al., 2019; Lou et al., 2019; Picco et al., 2021). Microsatellites are short tandem repeats of repetitive nucleotides that reside throughout the genome. Microsatellite stable (MSS) cells have two DNA repair mechanisms - (1) MMR machinery and (2) WRN to ensure genetic integrity. In MSS cells, disruption to either MMR or WRN does not lead to cell death. However, in MSI-high cells, MMR processes are compromised so the combination of inhibiting WRN leads to cell death due to this synthetic lethal relationship (Chan et al., 2019; Lieb et al., 2019; Picco et al., 2021). This WRN synthetic modality can be exploited for therapeutic value and has recently entered the clinic for the treatment of MSI-H cancers (NCT05838768 2018; NCT06004245 2023). A deeper understanding of DNA replication and repair regulatory mechanisms may provide insights that can be exploited for therapeutic-based purposes. Here, we present the identification of a new WRN degradation mechanism revealed by single molecule tracking (SMT). Upon WRN inhibition, WRN becomes trapped on chromatin and is SUMOylated by PIAS4. SUMOylated WRN is recognized by the ubiquitin E3 ligase RNF4, which then ubiquitinates WRN leading to p97/VCP-mediated chromatin extraction and ultimately proteasomal dependent degradation, providing new avenues for therapeutic targeting.

To investigate the consequence of WRN inhibition in WRN dynamics, we utilized a recently published clinical WRN inhibitor HRO761 (WRNi, Fig. 1a) (NCT05838768 2018) (Bordas et al., 2022). We first established a panel of MSI-H and MSS cells and tested for WRN sensitivity by depleting WRN using siRNAs (Extended Data Fig. 1a-d). Testing this compound in cells showed a >100-fold induction of DNA damage response in MSI-H cells compared to MSS cells, resulting in apoptosis and cell death (Fig. 1b-d, Extended Data Fig. 1g). Additionally, cellular toxicity was only observed in MSI-H, but not MSS cells, further highlighting the specificity of WRNi (Fig. 1e, Extended Data Fig. 1h). To profile the specificity of this compound *in vitro*, we purified WRN protein and its related homologue, BLM (Extended Data Fig. 1e-f). Using this recombinant system, we observed a 1000-fold difference in ATPase and helicase inhibition with WRN vs BLM (Fig. 1f-g, Extended Data Fig. 1i-j). Further highlighting the accumulation of DNA damage by WRNi, we observed an increase in nuclear size, indicative of DNA damage accumulation and potential cell senescence prior to apoptosis (Extended Data Fig. 2a-b) (Dedov et al., 2003; Rello-Varona et al., 2006; Kang et al., 2010). To enable imaging of proteins at the single molecule level we used CRISPR to generate endogenous HaloTag^TM^ WRN fusion protein (WRN^Halo^) in MSS (U2OS) and MSI-H (HCT-116) cells (McSwiggen et al., 2023). Prior to our imaging studies, we validated the successful tagging of WRN in these cells by depleting the HaloTag^TM^ WRN signal using a HaloTag^TM^ PROTAC (Extended Data Fig. 2c-d). Furthermore, comparison between WT and HaloTag^TM^-WRN HCT-116 cells showed no discernable differences between the two upon WRN inhibition (Extended Data Fig. 2e).

**Fig. 1:**
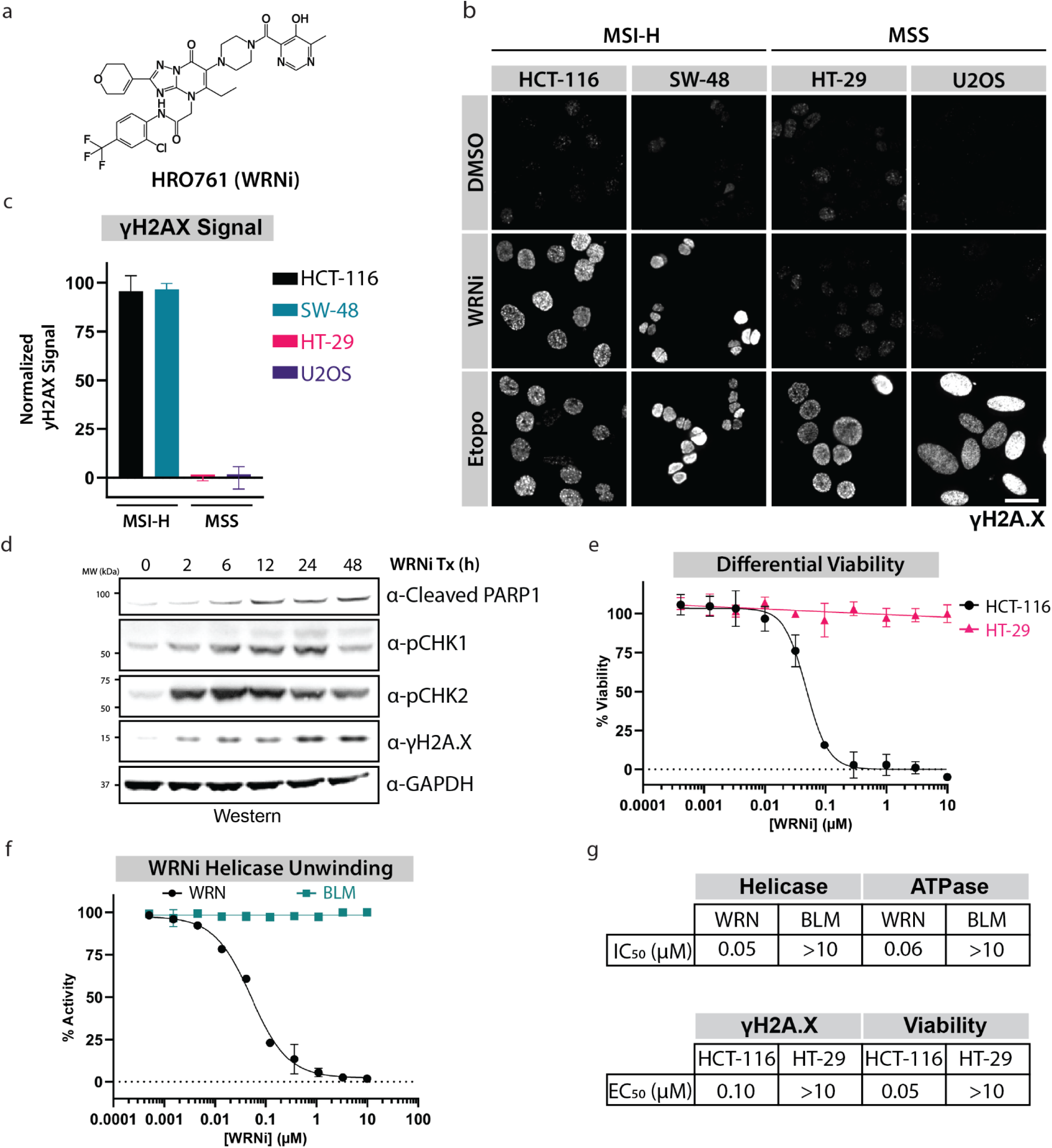
HRO761 is a specific and potent WRN inhibitor. **a.** Chemical structure of HRO761 (WRNi). **b.** WRN inhibition leads to induction of the DNA damage response in MSI-H cells. Phospho-histone H2A.X (Ser139) (γH2A.X) staining was used to visualize DNA damage in MSI-H (HCT-116, SW-48) and MSS (HT-29, U2OS) cells after 24 h treatment with indicated compounds. Scale bar = 20 μm. **c**. Quantification of γH2A.X signal levels in cells in **b**, normalized to DMSO and Etoposide. Graphs represent averages of at least 3 plate replicates, consisting of at least 6 fields of view (FOVs). **d**. WRN inhibition by WRNi leads to an induction of the DNA damage response resulting in apoptosis. HCT-116 cells treated with 10 μM WRNi and analyzed via Western to assess DNA damage response markers. **e**. Dose response curves measuring viability of HCT-116 cells or HT-29 cells after treatment with WRNi for 4 days. Graphs represent averages from *n* = 6 plates. **f**. Dose response curves measuring *in vitro* WRN or BLM unwinding activity after treatment with WRNi. Graphs represent averages from *n* = 4 replicates. WRN unwinding activity is normalized to DMSO and ATP-γ-S; BLM activity is normalized to DMSO and 10 μM BLMi. **g**. Summary of IC_50_ or EC_50_ values of WRNi in the indicated biochemical and cellular assays. All curve fits were done by fitting a 4-parameter logarithmic regression curve. All error bars represent standard deviation (s.d.). DMSO is dimethyl sulfoxide; BLMi is BLM inhibitor Compound 2; Etopo is etoposide. MW is molecular weight.

WRN undergoes a sub-compartmental translocation in response to DNA damage using standard immunofluorescence (Marciniak et al., 1998; von Kobbe and Bohr 2002). As previously reported, we observed robust translocation of WRN from the nucleolus to the nucleoplasm upon induction of DNA damage (Fig. 2a) (Kamath-Loeb et al., 2017; Veith et al., 2019; Zhu et al., 2021). We employed SMT to understand the link between WRN enzymatic activity mobility in cells using WRNi. Our fully automated SMT platform enables the measurement of thousands of experimental conditions and millions of cells per day (McSwiggen 2023). Recently we introduced substantial improvements to the platform, utilizing a light-sheet-based illumination strategy that improves both the throughput and quality of the SMT platform (Driouchi et al., submitted). Upon WRNi treatment, we observed a significant and dose-dependent decrease in average WRN mobility in the HCT-116 background (Fig. 2b-d, Extended Data Fig. 2f). Strikingly, no changes in protein diffusion coefficient were observed in U2OS, suggesting that the microsatellite state of the cell influenced the effect of WRNi on WRN dynamics (Fig. 2c, Extended Data Fig. 2g). Due to the virtue of SMT being a single molecule assay, we next sought to extract a more granular view of dynamic states of WRN in cells. To this end, we plotted the proportion of WRN proteins moving at a particular diffusion rate to generate a “state array” that provides a glimpse of the various states in which WRN is found, ranging from an immobile chromatin-bound state to a freely diffusing state (McSwiggen et al., 2023). We observed a decrease in the fastest-diffusing states and more than a 2-fold increase in the chromatin bound fraction of WRN in HCT-116 cells upon WRNi treatment (Fig. 2e-g). In contrast, we did not observe these changes in microsatellite stable U2OS cells or with treatment of general DNA damaging compounds (Fig. 2c and Extended Data Fig. 2g). The observed decrease in WRN diffusion coefficient was subsequently followed by a decrease in SMT spot density, suggesting possible degradation of the WRN protein (Fig. 2h).

**Fig. 2:**
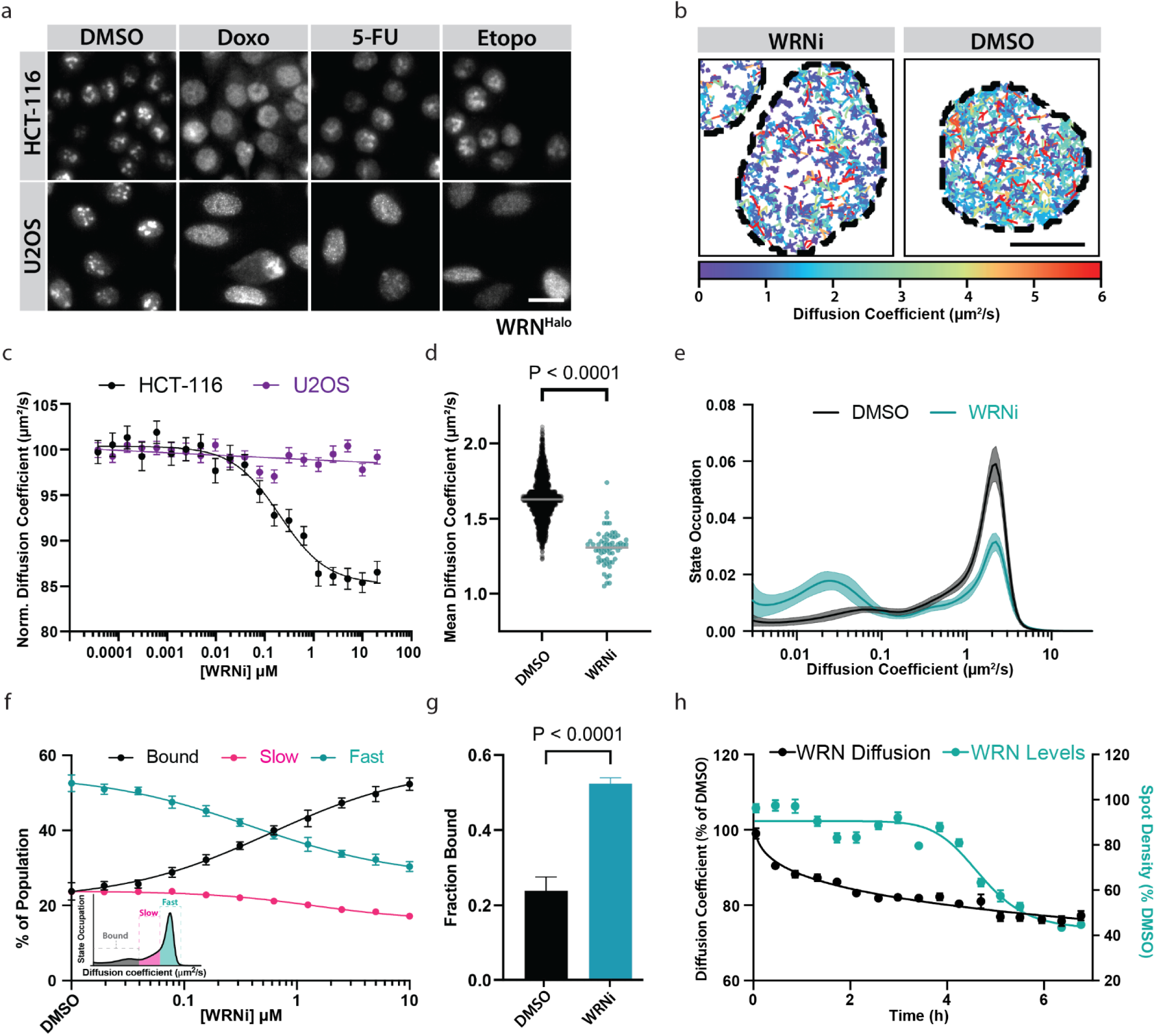
Single molecule tracking shows a change in WRN cellular dynamics in an MSI-H dependent manner. **a.** WRN^HALO^ cell lines, HCT-116 and U2OS, were validated by visualizing the subcellular localization of WRN in the presence or absence of DNA damaging compounds. WRN^Halo^ successfully translocates across compartments, suggesting a functional protein. Scale bar = 20 μm. **b**. The decrease in WRN diffusion coefficient upon inhibition can be captured by SMT, showing a dramatic slowdown of WRN protein. WRN inhibition leads to a decrease in molecule diffusion coefficient. Representative SMT track overlays over Hoechst nuclear stain outlines, in the presence or absence of 10 μM WRNi. Tracks are colored according to the diffusion coefficient of the molecules. **c**. Inhibition of WRN only affects its mean diffusion coefficient in MSI-H cells. Dose response curves with WRNi measuring the diffusion coefficient of WRN^HALO^ after 4 h treatments in HCT-116^WRN-HALO^ or U2OS^WRN-HALO^. Graph represents the average from *n =* 4 plates per condition. Error bars represent standard error of the mean (s.e.m.). All curve fits were done by fitting a 4- parameter logarithmic regression curve. **d**. Dot plot quantification of WRN diffusion coefficient from SMT movies after treatment with 10 μM WRNi. Each point represents the average WRN diffusion coefficient within all the nuclei in an FOV. *n* = 20 plates. Lines represent sample medians. **e**. WRNi shifts a large fraction of molecules from the free-diffusing state (“fast”) to the chromatin-bound (“bound”) state. Distribution of diffusive states in HCT-116^WRN-Halo^ cells showing the relative proportion of WRN molecules as a function of diffusion coefficient occupation, in the presence and absence of 10 μM WRNi. Shaded area represents s.d. **f**. Treatment with WRNi shows a dose-dependent increase in the bound fraction of WRN, suggesting binding of WRN onto chromatin. Dose response curves with WRNi measuring the different diffusive states of WRN protein. Inset is representation of how diffusive states are classified. Error bars represent s.d.. **g**. Quantification from **f** of the chromatin bound fraction of WRN^Halo^ protein in the presence or absence of 10 μM WRNi. Error bars represent s.d.. **h**. SMT can capture changes in protein diffusion before degradation is observed. Overlay of WRN^HALO^ diffusion coefficient and molecule spot densities after treatment with 10 μM WRNi over the indicated time points. Values are normalized to DMSO by dividing all FOV-level measurements at each time point by the median DMSO value at that time point, then multiplying by 100. *n* = 2 plates. Error bars represent s.e.m.. Curve fits were done by fitting a 4-parameter logarithmic regression curve. P-values were calculated using a two-tailed, unpaired t-test. DMSO is dimethyl sulfoxide; WRNi is HRO761; 5-FU is 5-fluorouracil, Doxo is doxorubicin.

We confirmed this observation by performing a WRNi time-course which resulted in a reduction of WRN protein levels in HCT-116 cells but not U2OS cells (Fig. 3a-b, Extended Data Fig. 3c). WRN degradation does not result from general DNA damage, but instead is specific to WRNi in a dose-dependent and proteasomal dependent way, as co-treatment of HCT-116 with the proteasome inhibitor carfilzomib rescues this effect (Extended Data Fig. 3a-b). Using cycloheximide, we determined that WRN inhibition reduced the half-life of WRN protein by over an order of magnitude, from 16.6h to 1.5h (Fig. 3b, and Extended Data Fig. 3d). The increased proportion of chromatin-bound (“bound”) WRN molecules, and the concomitant decrease in the freely diffusing (“fast”) population, shown in the state array data, suggests that inhibition of WRN by WRNi leads to its trapping on DNA (Fig. 2e-g). This mode of action has been widely studied with PARP1 and its inhibitors (Helleday 2011; Lord and Ashworth 2017).

**Fig. 3:**
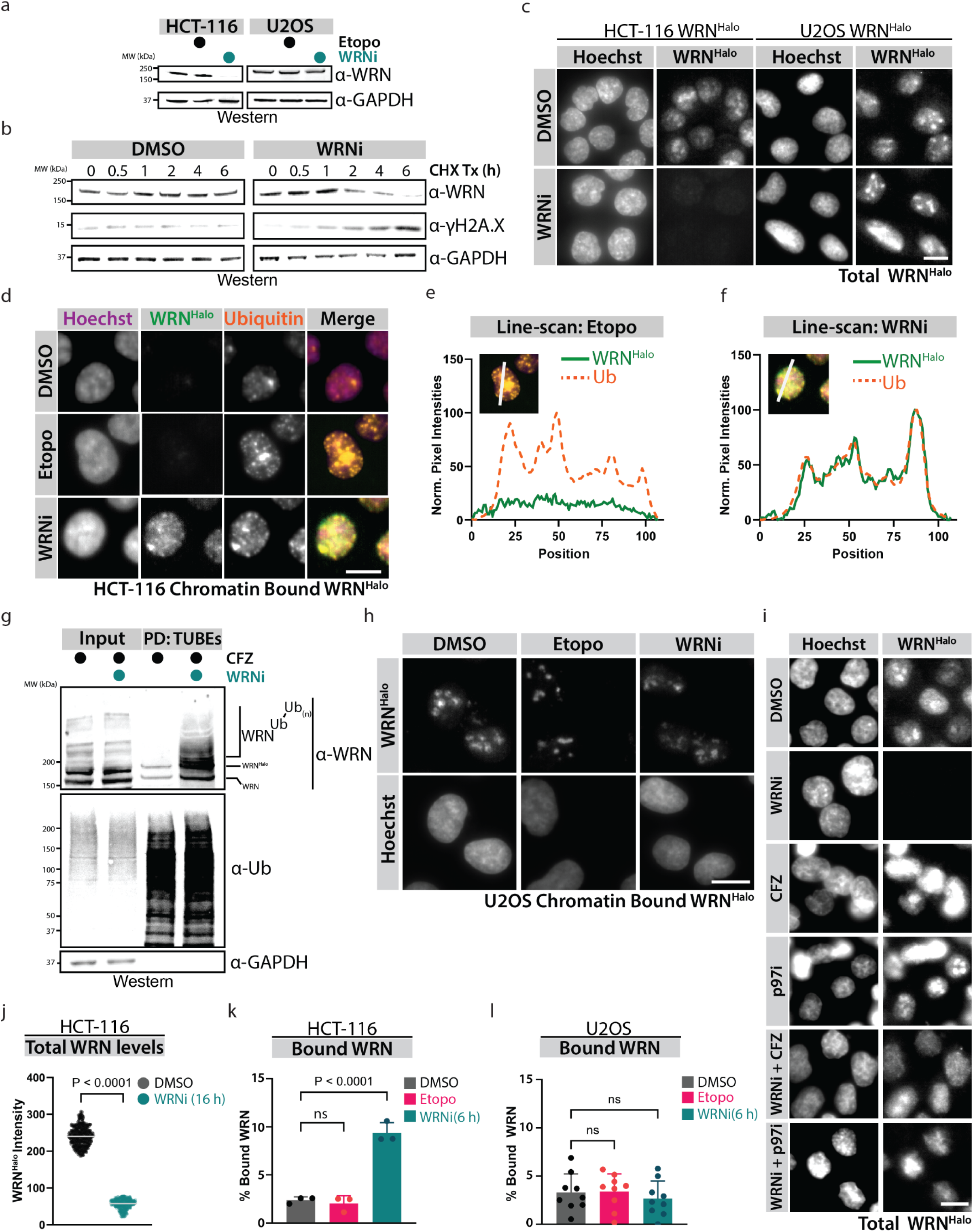
WRN Inhibition leads to its chromatin associated degradation. **a.** Inhibition of WRN leads to its degradation in an MSI-H dependent manner. HCT-116^WRN-Halo^ or U2OS^WRN-Halo^ were treated with 10 μM WRNi or etoposide for 16 h and analyzed by Western blot. **b**. Inhibition of WRN in MSI-H leads to a change in the half-life of WRN protein. HCT-116^WRN-Halo^ were treated with 100 μg/mL of cycloheximide (CHX) to inhibit protein synthesis in the presence or absence of 10 μM WRNi, then harvested at the indicated time points for Western blot analysis. Quantifications of CHX chase are in **Extended Data** Fig. 3d. **c**. Loss of WRN protein after treatment with WRNi can be visualized by microscopy. HCT-116^WRN-Halo^ or U2OS^WRN-Halo^ were treated with 10 μM WRNi for 16 h then imaged to visualize total WRN protein levels. Scale bar = 10 μm. **d**. WRN inhibition leads to WRN trapping on chromatin and its ubiquitylation. HCT-116^WRN-Halo^ cells were treated with 10 μM WRNi or etoposide for 6 h, at which point cells were permeabilized by treating with a mild detergent, then fixed and imaged. WRN signal retention is only seen when WRN is inhibited, and not with general DNA damaging compounds. Ubiquitin signal was also observed to colocalize with WRN^Halo^ when probing with a ubiquitin antibody. Scale bar = 10 μm. **e**. Line-scan quantification of etoposide treated cells from **d**, showing a lack of co-localization of the ubiquitin signal channel and the WRN signal channel. **f**. Line-scan quantification of WRNi treated cells from **d,** showing clear colocalization of the ubiquitin and WRN signal channels. **g**. WRN inhibition leads to its ubiquitylation. Tandem ubiquitin binding entities (TUBEs) pulldown (PD) of HCT-116 cells after treatment with 10 μM WRNi for 6 h in the presence of 1 μM carfilzomib (CFZ). Blotting for endogenous WRN using an α-WRN antibody shows higher molecular weight species in the TUBEs pull down only in the presence of WRNi. **h**. WRN chromatin trapping upon its inhibition is MSI-H dependent. Cells were treated with WRNi as in **d** but using the MSS cell line U2OS^WRN-Halo^. **i**. Degradation of WRN is dependent on the p97/VCP- proteasome axis. HCT-116^WRN-Halo^ were treated with 10 μM of WRNi and 1 μM of either CB-5083 (p97i) or CFZ for 6 h, then imaged. WRN protein degradation by WRNi is rescued upon co-treatment with CFZ or p97i. Quantifications are in **Extended Data** Fig. 3f. **j-l.** Quantifications of **c**, **d,** and **h.** Graph in **j** represents averages from *n* = 3 plates, with each individual point representing one well. Graph in **k** represents averages from *n* = 3 plates, each individual point is the average of 6 wells. Graph in **l** represents averages from *n* = 9 plates, each individual point is the average of 6 wells. Error bars represent s.d.. DMSO is dimethyl sulfoxide; WRNi is HRO761; CHX is cycloheximide; CFZ is carfilzomib; Etopo is etoposide. P-values were calculated using a two-tailed, unpaired Student’s t-test. ns = not significant. MW is molecular weight.

To investigate whether a similar mechanism of action was occurring upon WRN inhibition, we treated WRN^Halo^ cells with WRNi followed by permeabilization with a mild detergent to liberate soluble contents of the cells followed by fixation for immunolocalization studies (Illuzzi et al., 2022). These solubilization experiments revealed that treatment with WRNi led to the accumulation of WRN bound to chromatin based on the retention of WRN signal detected in the nucleus after treatment with detergent (Fig. 3d, 3k). Strikingly, we also observed increased colocalization of ubiquitin and WRN upon treatment with WRNi, further fortifying the link between WRN inhibition and the ubiquitin-proteasome pathway (Fig. 3d-f). This link was also validated by using tandem ubiquitin binding entities (TUBEs), which revealed a clear enrichment of higher molecular weight WRN species after WRNi treatment (Fig. 3g). Taken together, these data indicate that WRN is ubiquitylated upon its inhibition. Consistent with the lack of γH2A.X induction observed in U2OS cells upon WRN inhibition, WRNi treatment in this MSS background did not result in WRN trapping or degradation, suggesting that microsatellite instability plays a role in the regulation of WRN upon its inhibition (Fig. 3c, 3h, 3l and Extended Data Fig. 3e). Replication machinery has the potential to become stalled on DNA due to blockade by other bound proteins or DNA lesions (Edenberg et al., 2014; Liao et al., 2018; Le et al., 2023). To mitigate trapping, cells take advantage of ubiquitin-mediated proteasomal degradation in which these stalled proteins are marked for degradation by classical proteasome pathways after ubiquitin deposition (Vaz et al., 2013; Franz et al., 2016; Challa et al., 2021). p97/VCP has been implicated in the extraction of ubiquitylated membrane-bound, and chromatin-bound proteins. Therefore, we next investigated if the p97/VCP-proteasome axis was responsible for degradation of WRN (Rape et al., 2001; Jarosch et al., 2002; Wojcik et al., 2004; Meyer et al., 2012). Indeed, upon co-treatment with p97/VCP or proteasome inhibitors we were able to rescue the WRN degradation phenotype induced by WRNi in HCT-116 cells, suggesting an endogenous ubiquitin-dependent regulatory paradigm for WRN (Fig. 3i, Extended Data Fig. 3f). This finding was further validated using cellular SMT (Extended Data Fig. 3g) (Kim and Crews 2013; Anderson et al., 2015). Having determined that WRN degradation by WRNi treatment is mediated by the p97/VCP-proteasome axis, we next set out to identify the ubiquitin E3 ligase that activates this process. Depletion of E3 ligases, MIB1 and MDM2, that have been reported to regulate WRN did not prevent the degradation of WRN protein upon WRN inhibition (Extended Data Fig. 4a) (Liu et al., 2019; Li et al., 2020). Therefore, we set out to perform a ubiquitome-focused phenotypic siRNA screen and identified six potential genes that upon knock-down rescue WRNi-mediated degradation of WRN or regulate WRN protein levels in other ways (Fig. 4a and Extended Data Fig. 4b). Further validation of these siRNA hits identified RNF4 as the ligase responsible for WRN degradation upon WRNi treatment (Fig. 4a-b). Knock-down of RNF4 showed the most significant rescue of the phenotype, with all RNF4 siRNA oligos rescuing WRN protein levels to at least 75% of control treated cells (Extended Data Fig. 4c). We speculate that the partial rescues from the other ubiquitin modulators indicate putative genetic interactors, but the lack of a full rescue effect suggests these are not direct modulators, such as UBE2D3, which is a known E2 conjugating enzyme for RNF4 (Fig. 4c and Extended Data Fig. 4c) (DiBello et al., 2016; Roman-Trufero and Dillon 2022). We also observed a rescue of the WRN degradation phenotype using SMT by measuring the density of WRN molecules after RNF4 siRNA knock-down (Extended Data Fig. 4d). Depletion of RNF4 abolished the ubiquitylation of WRN after WRNi treatment, further establishing that RNF4 is responsible for WRN degradation (Fig. 4d-f and Extended Data Fig. 4e). Surprisingly, depletion of RNF4 only slightly rescued the change in WRN diffusion coefficient after inhibition by WRNi, suggesting that WRN remains trapped on chromatin in the absence of ubiquitylation (Fig. 4g). Trapped WRN is detrimental for genome integrity, and failure to remove and degrade trapped WRN results in further DNA damage, as evidenced by increased yH2A.X accumulation following knockdown of RNF4 (Ext data fig 4f). Co-treatment with the E1 ubiquitin activating enzyme inhibitor TAK-234 (E1i) and WRNi led to a slight decrease in the diffusion coefficient of WRN, which could suggest that inhibited WRN remains bound to chromatin if it cannot be ubiquitylated (Hyer et al., 2018) (Fig. 4h). State array analysis revealed that the global inhibition of ubiquitylation by E1i alone led to an increase in the slow diffusing fraction of WRN protein (Fig. 4i, j). This indicates that ubiquitylation is indeed required for the regulation of WRN beyond its function to drive protein degradation as has been shown for various cellular processes (Manford et al., 2020; Rodriguez-Perez et al., 2021; Padovani et al., 2022). However, the state array analysis also revealed that co-treatment with E1i and WRNi lead to a slight but significant increase in the chromatin bound fraction of WRN (Fig. 4i, k). Taken together, these data suggest that WRN inhibition in MSI-H cells leads to an increase in chromatin binding, followed by ubiquitylation and subsequent degradation. This ubiquitylation is necessary to remove WRN from chromatin, as exemplified by the increase in WRN bound to chromatin in the presence of an E1 inhibitor. RNF4 is known as a <u>S</u>UMO-targeted <u>ub</u>iquitin ligase (STUBL) and has been reported to regulate trapped chromatin proteins such as PARP1, DNMT1, and TOP1A (Sun et al., 2020; Liu et al., 2021; Krastev et al., 2022). To investigate if SUMOylation is required for the degradation of inhibited WRN, we depleted cells of the SUMO ligase PIAS4. The depletion of PIAS4 provided a modest, yet significant rescue of WRN degradation upon WRNi treatment (Fig. 5a, 5c). We posit that this failure to completely rescue the degradation phenotype is due to redundancies among the different PIAS proteins. To investigate this, we co-treated cells with the SUMO E1 activating enzyme inhibitor ML-792 (SUMOi) and WRNi, leading to a full rescue of the degradation phenotype by various means of detection, including SMT (Fig. 5b-f and Extended Data Fig. 6a-b) (He et al., 2017). Furthermore, upon SUMOylation inhibition, we failed to pull down ubiquitylated WRN even in the presence of WRNi (Fig. 5g), indicating that the SUMO cascade is necessary to initiate ubiquitylation of WRN. This is evident by the appearance of higher molecular weight WRN, which corresponds to ubiquitylated and SUMOylated WRN, as they disappear upon co-treatment with SUMOi (Fig. 5h). Co-treatment with SUMOi and WRNi lead to a further decrease in WRN diffusion coefficient (Fig. 5i). State array analysis indicated that this decrease was due to an increase in the “slow” fraction of WRN, further suggesting that the SUMO/ubiquitin cascade is important for the regulation of inhibited WRN (Fig. 5j). Indeed, inhibition of the SUMO-Ubiquitin-p97/VCP cascade lead to an increase in chromatin-bound WRN following inhibition by WRNi (Extended Fig. 5a-b). Taken together, these data clearly show that small molecule inhibition of WRN leads to its trapping onto chromatin, leading to its SUMOylation and subsequent ubiquitylation by RNF4 which ultimately results in proteasomal degradation (Fig. 5m). This work has elucidated a novel molecular pathway that regulates WRN activity upon its inhibition. The identification of WRN as a synthetic lethal target in MSI-H cancers has opened new avenues for therapeutic intervention of these malignancies. Clinical medicine has been successful in the selective killing of cancer cells mediated by DNA binding protein inhibition [etoposide citation]. For example, the enzyme, Poly ADP-ribose polymerase (PARP) can induce specific cell toxicity by PARP trapping. PARP carries out its enzymatic function by repairing single-stranded DNA breaks, but inhibition leads to trapping on DNA. As a result, PARP function is attenuated leading to an accumulation of DNA damage, stalled replication forks and cytotoxicity (Shen et al., 2015; Hopkins et al., 2019; Rose et al., 2020). Understanding the mechanistic underpinnings of WRN regulation has implications for treating MSI-H cancers in the clinic. WRN that cannot be released from chromatin after trapping is detrimental and leads to persistent DNA damage, even after removal of WRN inhibition (Fig. 5l and Extended Data Fig. 6c-d). This regulatory cascade has various points in which potential therapeutic interventions are possible, such as co-inhibition of the SUMOylation cascade or depletion of RNF4, which results in synergistic shift in the EC_50_ of WRNi (Fig. 5k and Extended Data Fig. 6e-h). Many other areas of potential intervention exist, such as disruption of the translocation of WRN from the nucleolus to the nucleoplasm, or perturbation of the molecular players that recognize chromatin-trapped WRN protein. This work provides deeper insights into the regulation of WRN protein activity upon small molecule inhibition that was gained in part through cutting-edge super resolution microscopy techniques like cellular SMT. Such detailed understanding will likely be critical in defining optimal clinical strategies as first-in-class WRN inhibitors undergo clinical testing.

**Fig. 4:**
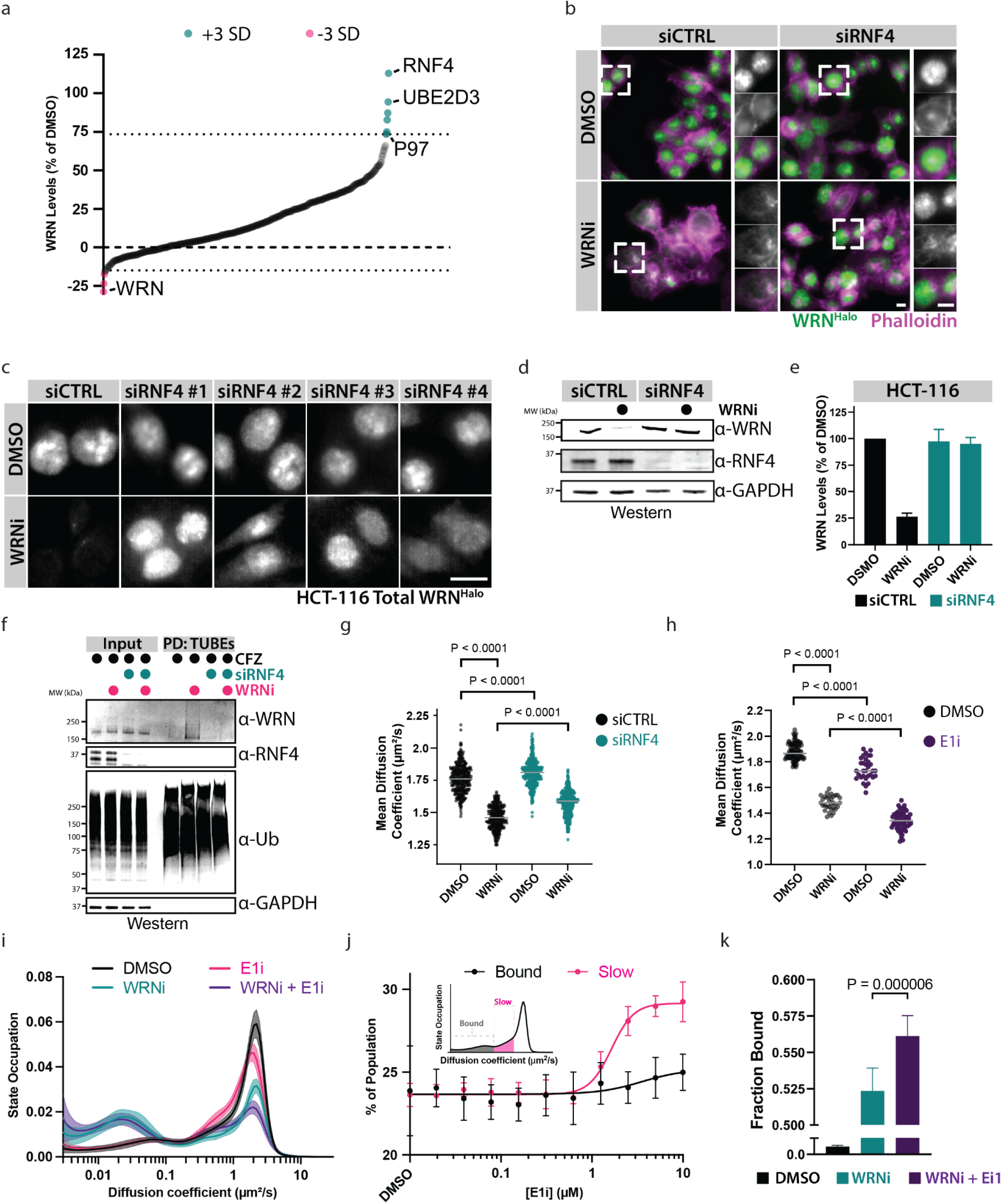
Phenotypic siRNA screen identified RNF4 as the ubiquitin E3 ligase targeting WRN for degradation. **a.** A phenotypic siRNA screen identified genes involved in the degradation of WRN after WRNi treatment. Colored circles indicate hits that are 3 standard deviations (SD) from the mean. Quantification of WRN protein levels after treatment with SMARTpool siRNAs in the presence or absence of 10 μM WRNi for 24 h. Depletion of RNF4 leads to stabilization of WRN protein in the presence of WRNi. UBE2D3 is an E2 known to interact with RNF4. **b**. Representative images of HCT-116^WRN-Halo^ cells the siRNA screen performed in **a,** showing a rescue of WRNi-induced degradation of WRN when RNF4 is depleted. Scale bars = 10 μm. **c**. Decomplexifying the siRNA SMARTpool validates RNF4 as the ligase responsible for WRN degradation by WRNi treatment. Representative images of HCT-116^WRN-Halo^ cells show all individual RNF4 siRNA oligos rescue the degradation phenotype induced by WRNi. Scale bar = 10 μm. Quantifications are in **Extended Data** Fig. 4c. **d**. Western blot analysis validates RNF4 as the ligase responsible for degradation of WRN induced by WRNi. HCT-116 cells were treated with siRNF4 oligos for 24 h, then treated with or without 10 μM WRNi for an additional 24 h, at which point cells were lysed and analyzed by Western blot with indicated antibodies. **e**. Quantifications of **d**. Graphs represent the mean value of *n* = 2 Western blot runs. Error bars represent s.d.. **f**. Depletion of RNF4 directly prevents ubiquitylation of WRN after WRNi treatment. HCT-116 cells were treated with siRNF4 oligos for 24 h, then cotreated with or without 10 μM of WRNi for 6 h. All samples were treated with CFZ to stabilize ubiquitin-modified proteins. TUBEs pulldowns (PD) and Western blot analysis were performed, probing with indicated antibodies. High molecular weight WRN species are detected in the TUBEs PD when treated with WRNi, which are not observed when treated with siRNF4 oligos, suggesting ubiquitylation of WRN is dependent on RNF4. **g**. RNF4 depletion leads to a slight but significant increase in the mean diffusion coefficient of WRN. Dot plots of WRN diffusion coefficient via SMT after co-treatment with siRNF4 and either DMSO or WRNi. Each point represents the average WRN diffusion coefficient within all the nuclei in an FOV. n = 4 plates. Lines represent sample medians. **h**. The ubiquitin pathway is involved in regulating WRN dynamics. Dot plots of WRN diffusion coefficient via SMT after co-treatment with the ubiquitin-activating enzyme (E1) inhibitor (E1i) and either DMSO or WRNi. E1i treatment caused a reduction in the diffusion coefficient of WRN. This decrease in diffusion was exacerbated by cotreatment with WRNi. Each point represents the average WRN diffusion coefficient within all the nuclei in an FOV. *n* = 4 plates. **i**. Distribution of diffusive states in HCT- 116^WRN-Halo^ cells showing the relative proportion of WRN molecules as a function of diffusion coefficient occupation after treatment with 1 μM E1i in the presence or absence of 10 μM WRNi. E1 inhibition leads to a shift from the “fast” to the “slow” state of WRN molecules. This shift is exacerbated by the cotreatment with WRNi. Shaded area represents s.d.. **j.** Treatment with E1i shows a dose-dependent increase in the slow fraction of WRN, suggesting that the ubiquitin pathway regulates WRN dynamics. Dose response curves with E1i measuring the “bound” and “slow” fractions of WRN protein. Inset is representation of how diffusive states are classified. Error bars represent s.d.. **k**. Bar graph quantification of the fraction bound of WRN^Halo^ protein after WRNi treatment in the presence or absence of E1i. Error bars represent s.d. DMSO is dimethyl sulfoxide; WRNi is HRO761; E1i is TAK-243; CFZ is carfilzomib. P- values were calculated using a two-tailed, unpaired Student’s t-test. ns = not significant. MW is molecular weight.

**Fig. 5:**
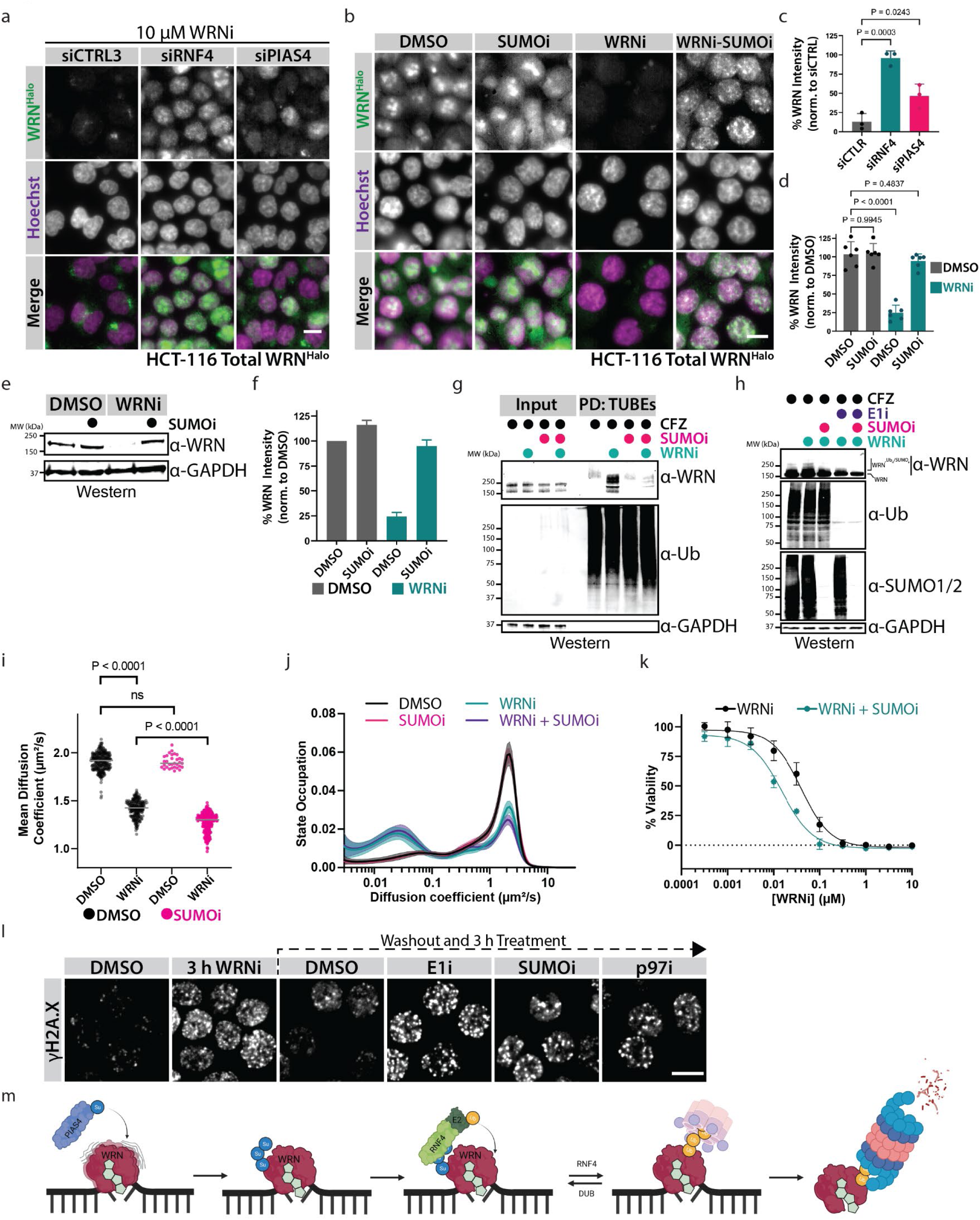
The PIAS4-RNF4 axis is responsible for the chromatin-associated degradation of WRN. **a.** WRN degradation is dependent on the SUMO E3 ligase PIAS4. HCT-116^WRN-Halo^ cells were treated with the indicated siRNA oligos for 24 h, and subsequently treated with 10 μM of WRNi for 24 h then imaged. WRN degradation is rescued by the depletion of RNF4, and partially rescued by PIAS4 depletion. **b**. SUMOylation is required for WRN degradation induced by WRNi. HCT-116^WRN-Halo^ cells were treated with 1 μM of the SUMO-activating enzyme (SAE) inhibitor ML792 (SUMOi) in the presence or absence of 10 μM WRNi, for 6 h and then fixed for imaging. Treatment with SUMOi results in a full rescue of the WRN degradation phenotype. **c** and **d**. Quantifications of **a** and **b,** respectively. Graphs represent the average of *n* = 3 plates for siRNA experiments and *n* = 6 plates for the small molecule experiments. Each dot represents the average of 6 wells, and 6 FOVs per well. Error bars represent s.d.. **e**. Whole cell lysate analysis shows SUMOylation is necessary for the degradation of WRN induced by WRNi. HCT-116 cells were treated with 1 μM SUMOi in the presence or absence 10 μM WRNi 16 h, at which point cells were lysed and analyzed by Western blot with indicated antibodies. Western blot analysis of cells showing WRN protein levels are stabilized when treated with SUMOi. **f.** Quantification of **e.** Graphs represent the mean value of *n* = 2 Western blot runs. Error bars represent s.d.. **g**. Inhibition of the SUMO pathway directly prevents ubiquitylation of WRN after WRNi treatment. HCT-116 cells were treated with 1 μM SUMOi in the presence or absence of 10 μM of WRNi for 6 h. All samples were treated with CFZ to stabilize ubiquitin-modified proteins. TUBEs pulldowns (PD) and Western blot analysis were performed, probing with indicated antibodies. High molecular weight WRN species are detected in the TUBEs PD when treated with WRNi, which are not observed when treated with SUMOi, suggesting that the SUMO pathway is required for the ubiquitylation of WRN upon its inhibition by WRNi. **h**. Whole cell lysate analysis by Western blot showing that higher molecular weight WRN species are dependent on the SUMO and ubiquitin pathways. Co-treatment with WRNi in the presence or absence of E1i and SUMOi show the disappearance of higher molecular weight species when co-treated with SUMOi or E1i, and a full disappearance of these species when co-treated with both SUMOi and E1i. **i**. Dot plots of WRN diffusion coefficient via SMT after co-treatment with SUMOi and either DMSO or WRNi. Each point represents the average diffusion coefficient of all the nuclei in an FOV. n = 4 plates. Lines represent sample medians; P-value was calculated using a two-tailed, unpaired t-test. Co-treatment with both compounds leads to a further decrease in the diffusion coefficient of WRN. **j**. Distribution of diffusive states for WRN^HALO^ in HCT-116^WRN-Halo^ after treatment with 1 μM of SUMOi in the presence or absence of 10 μM WRNi, showing that SUMOi leads to a decrease of the “fast” moving WRN molecules. **k**. Inhibition of the SUMO pathway synergizes with WRN inhibition. Dose response of WRNi in HCT-116 cells treated in the absence or presence of 10 nM SUMOi showing a shift in the EC_50_ viability of WRNi. **l**. Failure to extract trapped WRN leads to persistent DNA damage. Treating HCT-116 cells with WRNi for 3 h, then washing and replenishing with fresh growth media containing 1 μM of the indicated inhibitors show a retention in DNA damage as measured by γH2A.X when the SUMO-ubiquitin axis is perturbed. Scale bar = 10 μm. **m**. Model for targeting trapped WRN to proteasomal degradation by WRNi. WRN that is bound to chromatin surveying DNA damage in MSI-H cells becomes trapped upon inhibition by WRNi. This stalled WRN is SUMOylated by the SUMO ligase PIAS4. SUMOylated WRN recruits the STUbL RNF4, leading to its ubiquitylation. Ubiquitylated WRN is extracted from chromatin by p97/VCP, leading to its degradation by the proteasome. DMSO is dimethyl sulfoxide; WRNi is HRO761; E1i is TAK-243; SUMOi is ML-792; CFZ is carfilzomib. P-values were calculated using a two-tailed, unpaired Student’s t-test. ns = not significant. MW is molecular weight.

## Acknowledgements

The authors extend their deepest gratitude to all the employees and consultants of Eikon, past and present, especially, Pratir Doshi, Nick Vaquera, Jeff Dove, Tiffany Cheng, Roma Moore, Bruno da Rocha Azevedo, Madhu Mena, Emily Kirkeby, Puneet Kumar, Dave Piotrowski, Melissa Dumble, Geeta Sharma, and Alex Therian. Their tireless work enabled the experiments described here. We thank Roger Perlmutter, Robert Tjian for helpful discussions and critical feedback on the direction of our investigation and on the resulting manuscript. Eikon Therapeutics provided all funding.

## Materials and Methods

### Tissue Culture

HCT-116 (MSI genotype), U2OS, RKO, SW48, DLD-1 and HT-29 (MSS genotype) cells were used. Cells were cultured in Gibco McCoy’s 5A Medium (Thermo, 16600082) containing 10% fetal bovine serum (FBS), 1X GlutaMAX supplement (Thermo, 35050061), 1X MEM Non-Essential Amino Acids solution (Thermo, 11140076) and 50,000 Units/μg of Penicillin/Streptomycin (Thermo, 15140122) at 37°C and 5% CO_2_. Cells were seeded on Greiner 384-Well Black Microplate with Optical-Bottom (Greiner, 781906) using a Thermo Multidrop Combi. HCT-116 cells were seeded at 2,500 cells per well and HT-29 cells were seeded at 3,000 cells per well and incubated at 37°C and 5% CO_2_ for 24 hours before compound treatment. Compounds were added to the plates using a Beckman Echo 655 acoustic liquid handler. Compound treated plates were incubated for 24 h at 37°C and 5% CO_2_.

### Cell Line Engineering

WRN^Halo^ HCT-116 and U2OS were generated by nucleofection of ribonucleoprotein (RNP) complexes (DeWitt et al., 2017) using a guide (5’ – AAAGATGAGTGAAAAAAAAT – 3’) targeting the N-terminus of the *WRN* gene locus and a megamer coding for the Halo tag as a donor template (5’ – TATTGTATCTGTTTTGTTTTGTGATTCTAGCTCTTATAACCTATGCTTGGACCTAGGTGTCATAACTTACTTTAAATATGTATGTTTGGTTTTCATTCATATTGACAGTACTACCTCTCAGTTTTCTTTCAGATATTGTTTTGTATTTACCCATGAAGACATTGTTTTTTGGACTCTGCAAATACCACATTTCAAAGATGGCAGAAATCGGTACTGGCTTTCCATTCGACCCCCATTATGTGGAAGTCCTGGGCGAGCGCATGCACTACGTCGATGTTGGTCCGCGCGATGGCACCCCTGTGCTGTTCCTGCACGGTAACCCGACCTCCTCCTACGTGTGGCGCAACATCATCCCGCATGTTGCACCGACCCATCGCTGCATTGCTCCAGACCTGATCGGTATGGGCAAATCCGACAAACCAGACCTGGGTTATTTCTTCGACGACCACGTCCGCTTCATGGATGCCTTCATCGAAGCCCTGGGTCTGGAAGAGGTCGTCCTGGTCATTCACGACTGGGGCTCCGCTCTGGGTTTCCACTGGGCCAAGCGCAATCCAGAGCGCGTCAAAGGTATTGCATTTATGGAGTTCATCCGCCCTATCCCGACCTGGGACGAATGGCCAGAATTTGCCCGCGAGACCTTCCAGGCCTTCCGCACCACCGACGTCGGCCGCAAGCTGATCATCGATCA GAACGTTTTTATCGAGGGTACGCTGCCGATGGGTGTCGTCCGCCCGCTGACTGAAGTCGAGATGGACCATTACCGCGAGCCGTTCCTGAATCCTGTTGACCGCGAGCCACTGTGGCGCTTCCCAAACGAGCTGCCAATCGCCGGTGAGCCAGCGAACATCGTCGCGCTGGTCGAAGAATACATGGACTGGCTGCACCAGTCCCCTGTCCCGAAGCTGCTGTTCTGGGGCACCCCAGGCGTTCTGATCCCACCGGCCGAAGCCGCTCGCCTGGCCAAAAGCCTGCCTAACTGCAAGGCTGTGGACATCGGCCCGGGTCTGAATCTGCTGCAAGAAGACAACCCGGACCTGATCGGCAGCGAGATCGCGCGCTGGCTGTCGACGCTCGAGATTTCCGGCGAAAACCTGTATTTTCAGAGCAGTGAAAAAAAATTGGAAACAACTGCACAGCAGCGGAAATGTCCTGAATGGATGAATGTGCAGAATAAAAGATGTGCTGTAGAAGAAAGAAAGGTATGTTGTTCATTGACTATTCTTTTGGGTGAGAAATTTAATTTATATTTGACTGTGCAAAGAGTCAGTTGTTACTTGTAAACTTCAAGTCATTGTTTAGGTCAGAG – 3’). RNPs were assembled with Alt-R™ S.p. HiFi Cas9 Nuclease V3 (IDT, 1081060) in a 10 μl reaction of 1 μM of Cas9, 120 pmol sgRNA, and 100 pmol ssODN in Cas9 buffer (20 mM HEPES 7.5, 150 mM KCl, 10% glycerol, 1 mM TCEP). Reactions were gently mixed for 30s and incubated for 10min at room temperature. RNP complexes and 200k cells resuspended in 20 μl buffer SE (Lonza) were added to a nucleofection strip and the mixture pulsed with program EH-100 (Lonza 4D-Nucleofector). Cells were plated into 96-well plates for recovery with a fresh media change after 1 day. Two days after RNP electroporation, pooled cells then were single-cell seeded into 384-well plates. After single-cell derived clones emerged, they were split into 2 plates, one of which was used to detect positive clones containing the desired HaloTag^TM^ sequence through Sanger sequencing.

### Antibodies and Reagents

Goat anti-Rabbit IgG Alexa Fluor Plus 647 (Thermo, A32733), anti-phospho-Histone H2A.X (Ser139)-clone JBW301 (γH2A.X, Millipore, 05-636-I), anti-WRN clone EPR6392 (abcam, ab124673), Goat anti-Rabbit IgG (H+L)-HRP (Invitrogen, 31460), anti-GAPDH-HRP (CST, 31460), anti-Ubiquitin, clone P4D1 (for WB; CST, 3936), anti-Ubiquitin, clone FK2 (For IF; Millipore sigma, 04-263), anti-SUMO-2/3, clone 18H8 (CST, 4971S), anti-phospho-CHK1 (Ser345; CST, 2341), phospho-Chk2 (Thr68; clone C13C1; CST, 2661), cleaved PARP1 (Asp214; clone D64E10; CST, 5625), anti-RNF4 (R&D Systems, AF7964), anti-MDM2 (R&D Systems, D1V2Z), anti-MIB1 (CST, 4400).

JF549 (Promega, GA1110), etoposide (CST, 2200), 5-FU (5-fluorouracil; Selleckckem, S1209), doxorubicin (CST, 5927), HaloPROTAC3 (Promega, GA3110), HaloPROTAC-E (MedChem Express, HY- 145752), MLN-4924 (CST, 85923), carfilzomib (CST, 15022), TAK-243 (MedChem Express, HY-100487), ML-792 (MedChem Express, HY-108702).

### Immunofluorescence and DNA Damage Quantification

Cell culture medium was removed using the Blue Cat Bio Blue®Washer. Paraformaldehyde solution (8% PFA (Electron Microscopy Sciences, 15710-S), 1X phosphate buffered solution (PBS) (Teknova P0191)) was added to the cells via Thermo Multidrop® Combi and treated for 7 min at room temperature to fixate the cells. The paraformaldehyde solution was removed with the Blue Cat Bio BlueWasher. Triton solution (0.1% TritonX-100 (Sigma, T8787), 1X PBS) was added to the cells via Thermo Multidrop Combi and treated for 15 min at room temperature to permeabilize the cells. The triton solution was removed with the Blue Cat Bio BlueWasher. Serum solution (10% Goat Serum (Gibco, 16210), 1X PBS) was added to the cells via Thermo Multidrop Combi and treated for 15 min at room temperature to block the cells. The serum solution was removed with the Blue Cat Bio BlueWasher. Primary antibody solution (1:1000 primary antibody, 10% Goat Serum, 1X PBS) was added to the cells via a Thermo Multidrop Combi and treated for 3 hours at room temperature with shaking. The primary antibody solution was removed with the Blue Cat Bio BlueWasher. Secondary antibody solution (secondary antibody (1:1000), Hoechst 33342 (AnaSpec, AS-83210) (1:3000), 10% Goat Serum, 1X PBS) was added to the cells via Thermo Multidrop Combi and treated for 30 min at room temperature, covered from light, with shaking. Cells were washed with PBS 3 times using the Blue Cat Bio BlueWasher with a final dispense of PBS. Sealed plates were imaged using an ImageXpress Micro slit confocal microscope (Molecular Devices) using a 40x water immersion objective, and 6 fields of view per well. Exposure parameters were optimized to prevent pixel saturation for each channel. Images were analyzed using MetaXpress Custom Module Editor, by using a Hoechst mask to identify nuclei, then measuring the integrated AlexaFluor 648 intensity across all nuclei in the field of view (FOV) and then averaged. Percent γH2A.X signal was calculated by using the following equation: %S = (T - C_pos_)/(C_pos_ – C_neg_) * 100, where %S is percent γH2A.X signal and T is the measured γH2A.X fluorescence of the wells treated with test compound. The effective compound concentration leading to a 50% induction of γH2A.X signal (EC_50_), and the resulting cell γH2A.X signal measured at the highest tested compound tested (c_max_) was carried out by fitting a 4-parameter non-linear regression using GraphPad Prism. At least three biological replicates were done per compound tested.

### Western Blotting

Samples were lysed in 1x NuPAGE LDS sample buffer (Invitrogen, NP0008), sonicated, and heated at 75 °C for 10 min. Samples were normalized to protein concentration and volume using a Pierce 660 reagent kit (Invitrogen, 22660). SDS-Page was performed using standard protocols (ref). Western blot transfers were performed using an iBlot2 nitrocellulose membrane system (Invitrogen, IB23001). Membranes were blocked with Pierce™ Protein-Free Blocking Buffer (Thermo Scientific, 37572) for 30 min at room temperature, and subsequently probed with desired antibodies.

### Cell Viability Assay

To seed cells, cells were trypsinized and resuspended in complete culture media to the desired concentration (HCT-116: 10,000 cells/mL; RKO: 15,000 cells/mL; SW-48, HT-29, and SW-480: 25,000 cells/mL; U2OS: 7,000 cells/mL). Cell suspensions were seeded in 50 μL of complete culture media and plated onto 384-well white clear-bottom plates (ThermoFisher Cat# 164610) using a Multidrop^TM^ Combi liquid dispenser in the slowest setting in triplicate. After 24 h, cells were treated with compounds in a 10- pt, 3.16 step serial dilution using an Echo acoustic liquid dispenser (Beckman). DMSO was backfilled to a final concentration of 0.1%. 10 μM etoposide (C_pos_) and DMSO (C_neg_) were used as reference compounds. 96 h post compound addition, plates were evacuated to decant all media. To measure viability, cellular ATP concentrations were measured by adding 20 μL of 1x CellTiterGlo 2.0 (CTG) solution (1 part PBS, 1 part CellTiterGlo2.0 stock solution; Promega Cat# G9241) using a MultidropTM Combi liquid dispenser and measuring the luminescence on a SpectraMax iD3 (Molecular Devices).

Percent viability was calculated by the following equation: %V = (T - C_pos_)/(C_pos_ – C_neg_) * 100, where %V is percent viability and T is the measured luminescence of the wells treated with test compound. The effective compound concentration leading to a 50% reduction in viability (EC_50_), and the resulting cell viability measured at the highest tested compound tested (c_max_) was carried out by fitting a 4-parameter non-linear regression using GraphPad Prism. At least three biological replicates were done per compound tested.

### Tandem Ubiquitin Binding Entities (TUBE) Pull Down

HCT-116 cells were grown as described above. 10^6^ cells were seeded on 10 cm tissue culture dishes (ThermoFisher, 150464), and allowed to attach for 48h. Cells were subsequently treated with compounds for 8 h ([ML-792] = 1 μM, [Carfilzomib] = 10 nM, [TAK-243] = 10 nM, [WRNi] = 10 μM), harvested by trypsinization, centrifuged (300 x g), and flash frozen in liquid nitrogen. Cell pellets were lysed in 500 μL of TUBE lysis buffer (150 mM NaCl, 2 mM MgCl_2_, 25 mM HEPES pH 7.0, 0.03% SDS, 1 M urea, 10 μM PR-619, 1x protease inhibitor cocktail) for 10 min on ice, with frequent mixing. While cell lysis was occurring, 100 μL of slurry/reaction of TUBE magnetic beads (UM501M, LifeSensors) were washed in 1x PBS three times, and once with TUBE lysis buffer. After cell lysis, lysates were cleared (20,000 x g, 10 min), inputs were taken, and lysates were subsequently added to TUBE magnetic beads. Lysate-TUBE bead mixtures were incubated at 4 °C on a nutator for 2 h. Samples were subsequently washed five times with TUBE wash buffer (150 mM NaCl, 2 mM MgCl_2_, 25 mM HEPES pH 7.0, 0.5% Triton X-100, 1 M urea), and resuspended in 1X LDS buffer to prepare for downstream Western blot analysis.

### HaloTag^TM^ Protein labeling

Cells were seeded in black 384-well plates using a combidrop multidrop dispenser and seeded for at least 24 h in complete growth media at either 45K cells/mL or 100K cells/mL, depending on the experiment. To label HaloTagged proteins, JF549 (Promega, GA1110) was added to cells using an Echo acoustic liquid dispenser to a final concentration of 25 pM. Cells were incubated at 37 °C for 2 h, and subsequently washed 5 times in PBS, with the final wash leaving the wells empty. After washing, cells were either fixed in 4% paraformaldehyde (Electron Microscopy Sciences, 15710), or growth media was replenished to perform subsequent compound treatments.

### WRN Imaging Degradation Assays

Cell lines and growth conditions are identical to the above conditions. To seed cells for the assay, cells were trypsinized and resuspended in complete culture media to the desired concentration (WRN^Halo^ HCT-116: 50,000 cells, WRN^Halo^ U2OS: 30,000 cells/mL). Cell suspensions were seeded in 50 μL of complete culture media and onto 384-well black clear-bottom optical plastic plates (Greiner Bio-One Cat# 781097) using a Multidrop^TM^ Combi liquid dispenser in the slowest setting in triplicate. After 24 h, cells were labeled with Halo dye as described above and subsequently treated with compounds in a 10-pt, 3.16 step serial dilution using an Echo acoustic liquid dispenser and incubated at 37 °C in a humidified 5% CO_2_ incubator. 10 μM Halo-PROTAC3 (C_pos;_ Promega, GA3110) and DMSO (C_neg_) were used as reference compounds.24 h after compound treatments, cells were fixed in 4% paraformaldehyde for 10 min, washed three times with PBS, then blocked and permeabilized in PBS containing 10% goat serum and 0.1% triton X-100 containing the DNA counterstain Hoechst for 30 min. Plates were washed 3 times in PBS, and sealed with thermal foil seals.

Sealed plates were imaged using an ImageXpress Micro slit confocal microscope (Molecular Devices) using a 40x water immersion objective, and 6 fields of views per well. Exposure parameters were optimized to prevent pixel saturation for each channel. Images were analyzed using MetaXpress Custom Module Editor, by using a Hoechst mask to identify nuclei, then measuring the average Halo dye intensity across all nuclei in the FOV, averaged, and then background subtracted.

Percent WRN^Halo^ signal was calculated by using the following equation: %S = (T - C_pos_)/(C_pos_ – C_neg_) * 100, where %S is percent WRN^Halo^ signal and T is the measured WRN^Halo^ fluorescence of the wells treated with test compound. The effective compound concentration leading to a 50% induction of WRN^Halo^ signal (EC_50_), and the resulting cell WRN^Halo^ signal measured at the highest tested compound tested (C_max_) was carried out by fitting a 4-parameter non-linear regression using GraphPad Prism. At least three biological replicates were done per compound tested.

### Phenotypic Screening and Imaging

An arrayed human ON-TARGETplus ubiquitome siRNA SMARTpool library (Horizon Discovery, 106205-E2-01) was resuspended to a final concentration of 20 uM in 1x siRNA Buffer (Horizon Discovery, B-002000-UB-100). siRNA oligos were transferred onto black 384 μClear plates using an Echo acoustic dispenser to yield a final oligo concentration of 20 nM. 5 μL of opti-MEM (Gibco, 31985062) was added to resuspend transferred oligos. RNA oligo:lipid complexes were formed by adding 0.3 μL of Lipofectamine RNAiMAX (Invitrogen, 13778075) in 5 μL of opti-MEM. Complexes were incubated for 5 min before dispensing 50 μL of HCT-116 cells (100K cells/mL) and incubating in growth conditions described above. After 24 h, growth media was exchanged, and labeled with HaloTag^TM^ dye as described above. WRNi, or DMSO, was added at a final concentration of 10 μM, treated for 24 h, and subsequently fixed with 4% PFA. Cells were permeabilized with 0.1% triton-x 100 for 20 min to remove non-specific dye staining and imaged as described above.

### siRNA Depletions

Cells were processed as described for quantifying WRN levels after siRNA knock-down. For Western blot analysis, HCT-116 (90K cells/well) or U2OS (60K cells/well) cells were seeded in 6-well TC treated plates (Corning, 3516). siRNA depletions were performed following the recommended protocol for Lipofectamine RNAiMax. Samples were harvested in 1x LDS sample buffer and prepared for Western blot analysis as described.

### WRN In-Situ Trapping Assay

WRN^Halo^ Cells (HCT-116 or U2OS) were seeded in black 384-well plates using a combidrop multidrop dispenser and seeded for 48 h in complete media at 100K cells/mL (HCT-116) or 30 K cells/mL (U2OS). Cells were labeled with HaloTag^TM^ Dye and subsequently treated with desired compounds as described above for 8 h ([ML-792] = 1 μM, [Carfilzomib] = 10 nM, [TAK-243] = 10 nM, [WRNi] = 10 μM). Following compound treatments, samples were decanted to remove all media and treated with ice cold CSK buffer (25 mM PIPES pH 7.0, 300 mM NaCl, 2 mM MgCl_2_, 0.3 % TritonX-100, 200 mM sucrose) for 2 min, on ice. Without removing buffer in wells, cells were fixed in 4% PFA supplemented with Hoechst for 15 min and washed with 1x PBS 5 times using an AquaMax Plate washer (Molecular Devices). Samples were then imaged as described above. Quantifications were done by measuring the total intensity of the Halo dye signal and using the Hoechst channel as a nuclear mask.

### Protein Purification

Human WRN residues 480–1251 were cloned into pFastBac vector containing an 8xHis N-terminal tag and expressed in Sf9 insect cells. Harvested cells were resuspended in lysis buffer (50 mM HEPES, pH 7.5, 500 mM NaCl, 25 mM imidazole, 1mM TCEP, 5 U/mL benzonase (Millipore Sigma), EDTA-free cOmplete protease inhibitor cocktail tablet (Roche)) and lysed by addition of Insect PopCulture reagent (Millipore Sigma). Lysate was loaded on to a HiTrap TALON Crude column (Cytiva) and protein was eluted using 500mM Imidazole. Fractions containing WRN protein were collected, pooled, then diluted to in buffer to drop the NaCl concentration to ∼100mM. The pooled and diluted sample was then loaded onto a heparin column (Cytiva) and eluted using a step-wise gradient from 200mM to 1000mM NaCl. Fractions containing purified WRN protein were pooled, and buffer exchanged into 50mM HEPES, 150mM NaCl, 10% glycerol, 1mM TCEP using a HiPrep desalting column (Cytiva).

Human BLM residues 636-1298 were cloned into pFastBac vector containing an 8xHis N-terminal tag and expressed in Sf9 insect cells. Harvested cells were resuspended in lysis buffer (50 mM HEPES, pH 8, 200 mM NaCl, 0.5mM TCEP, 5 U/mL benzonase (Millipore Sigma), EDTA-free cOmplete protease inhibitor cocktail tablet (Roche)) and lysed by addition of Insect PopCulture reagent (Millipore Sigma). Lysate was loaded onto a Ni-NTA column (Thermo Fisher) and protein was eluted using 300mM Imidazole. Fractions containing BLM protein were collected, pooled, and diluted in buffer to drop the NaCl concentration to ∼50mM. The pooled and diluted sample was then loaded on a HiTrap Heparin column (Cytiva) and eluted with a linear gradient from 50mM to 1M NaCl. The pure fractions were pooled and concentrated and purified by size exclusion chromatography using S200 Increase (Cytiva) in 50mM HEPES pH 8, 200mM NaCl, 5% glycerol, 0.5mM TCEP.

### Helicase Unwinding and ATPase Assay

The helicase unwinding and ATPase were carried out in a multiplexed fashion based on previously published BLM assay (Chen et al., 2021). Single-stranded DNA was purchased from IDT: A: 5’-Cy3-GAACGAACACATCGGGTACGTTTTTTTTTTTTTTTTTTTTTTTTTTTTTT B: 5’-TTTTTTTTTTTTTTTTTTTTTTTTTTTTTTCGTACCCGATGTGTTCGTTC-IowaBlackFQ-3’ B-dark: 5’-CGTACCCGATGTGTTCGTTCY-3’ Strands A and B were annealed in TE + 50mM NaCl in a slowly-cooling thermocycler.

First, compounds in DMSO were dispensed to a 384-well white ProxiPlate (Perkin Elmer) using an Echo acoustic liquid handler. WRN or BLM Protein was diluted into assay buffer and 2.5 µL was dispensed into each well. Protein and compound then incubated at room temperature for 15 min. Following pre-incubation, 2.5µL of DNA and ATP in assay buffer was dispensed into each well to initiate the reaction and incubated for 20 min at room temperature. At the 20 minute time point, Cy3 fluorescence was read on an Envision plate reader to measure unwinding activity. ATP hydrolysis was then measured using ADP-Glo kit (Promega) by adding 5uL ADP-Glo reagent for 40 minutes followed by 10uL Kinase Detection Reagent for 1hour. Luminescence was measured using Envision plate reader. Data was normalized to DMSO (100% activity). The final reaction conditions are: 1mM ATP, 15nM dsDNA substrate, 1.5uM B-dark in reaction buffer composed of 50mM Tris-HCl pH 8.0, 50mM NaCl, 2mM MgCl2, 0.01% Tween-20, 2.5µg/mL poly(dI-dC), 1mM DTT, 1% DMSO, and 12.5nM WRN or 2.5nM BLM.

### Cellular SMT Sample preparation

For SMT experiments, WRN Halo-tagged HCT-116 cells were seeded in FluoroBrite DMEM (Thermo Fisher, cat. no. 1896701) supplemented with 10% FBS (Corning), 1% Penicillin/Streptomycin (Gibco, 1510-122), and 1% GlutaMAX (Gibco, 35050-061) on plasma-coated 384-well glass-bottom plates (Cellvis, P384-1.5H-N) at 1.5x10^4^ cells per well. WRN Halo-tagged U2OS cells were seeded in GlutaMAX-supplemented DMEM (Gibco, 10566-016) with 10% FBS and 1% Penicillin/Streptomycin at 6x10^3^ cells per well. Prior to treatment and imaging, HCT-116 cells were incubated at 37 °C and 5% CO_2_ for ∼48 hours to allow for cell adherence to the plates, while U2OS cells were incubated under the same conditions for 24 hours. For all SMT experiments, cells were treated with 10-40 pM of JF549-HTL (synthesized in-house) and 200 nM Hoechst 33342 (Thermo Fisher, cat. No. 62249) for 1 hour in complete medium at 37 °C and 5% CO_2_. Cells were then washed three times in PBS and twice in FluoroBrite DMEM supplemented with 2% FBS, 1% Penicillin/Streptomycin, and 1% GlutaMAX. All compounds were prepared on Echo Qualified 384- Well Low Dead Volume Source Microplates (Labcyte, cat. No. LP-0200) in DMSO and administered onto cells at a final 1:500 dilution in cell culture medium. Unless otherwise specified, cells were incubated with compounds for 4 hours at 37 °C prior to image acquisition. When possible, well replicate conditions were randomized across each plate. For all experiments, control conditions included vehicle (DMSO) treatment and wells lacking JF549-HTL to assess possible effects from detection of non-dye signal.

### Cellular SMT Image Acquisition

Unless otherwise stated, all image acquisition for cSMT was performed on a customized Nikon Eclipse Ti2-E inverted fluorescence microscope with a motorized stage. The microscope system was outfitted with a stage top environmental chamber with temperature and CO_2_ control (OKO labs), Nikon objective water dispenser, an Oblique Line Scanning (OLS) illumination module (Driouchi et al., submitted) with laser launch containing 405 nm, 560 nm, and 642 nm lasers, three-band emission filter set (ET 445/58m, FF01-585/40-25, FF01-676/37-25, Chroma), motorized filter wheel (Lambda 10-B; Sutter Instruments), and a high-speed sCMOS camera equipped with light-sheet mode capability (ORCA-Fusion BT, Hamamatsu). Images were acquired with a 60X 1.27 NA water immersion objective (CFI SR Plan Apo IR 60XC WI, Nikon, Japan). The environmental chamber was maintained at 37° C, 95% humidity, and 5% CO_2_. For each field of view, 150 SMT frames were collected at a frame rate of 100 Hz with a 407-microsecond stroboscopic laser pulse, and 1 frame in the Hoechst channel was subsequently collected for downstream registration of trajectories to nuclei. Each frame captured an FOV that was 1728 x 2304 pixels (187.14 x 249.52 microns) in size. Automated microscope control and image acquisition was performed using customized scripts in MicroManager.

For all experiments, conditions were tested by acquiring JF549 movies and Hoechst images at 4 FOVs per well, a minimum of 2 well replicates per plate, and a minimum of 2 plate replicates. Unless otherwise stated, reported averages for each condition are the mean value of all FOVs collected. For time course experiments, reported averages at each time point are the mean value of all FOVs collected across 6 consecutive wells per plate replicate.

### Cellular SMT Image Processing

Image acquisition yielded one JF549 movie and one Hoechst movie per FOV. The JF549 movie was used to track the motion of individual JF549 molecules, while the Hoechst movie was used for nuclear segmentation.

For tracking, we used a custom pipeline that operates in three sequential steps. First, dye molecules are detected using a generalized log likelihood ratio detector (Serge et al., 2008). The position of each detected emitter is then estimated using a Levenberg-Marquardt fitting routine (Levenberg 1944; Marquardt 1963; Laurence and Chromy 2010) with an integrated 2D Gaussian spot model (Smith et al., 2010) starting from an initial guess afforded by the radial symmetry method (Parthasarathy 2012). Detected emitters were then linked into trajectories using a custom algorithm. Briefly, this method first estimates the marginal probabilities of each potential link between particles using the graphical softmax operator (Cuturi 2013; Mena et al., 2018) applied to a Brownian motion model, then attempts to find the set of trajectories with maximum marginal log probability using a modification of Sbalzerini’s hill-climbing algorithm (Sbalzarini and Koumoutsakos 2005). Identical tracking settings were used for all movies in this manuscript.

For nuclear segmentation, all frames of the Hoechst movie were averaged to generate a mean projection. This mean projection was then segmented with a UNET-based convolutional neural network trained on human-labeled nuclei (Ronneberger et al., 2015). Each spot was then assigned to at most one nucleus using its subpixel coordinates. To recover dynamical information from trajectories, we used state arrays (Heckert et al., 2022), a Bayesian inference approach, with the “RBME” likelihood function and a grid of 100 diffusion coefficients from 0.01 to 100.0 µm^2^/sec and 31 localization error magnitudes from 0.02 µm to 0.08 µm (1D RMSD). After inference, localization error was marginalized out to yield a one-dimensional distribution over the diffusion coefficient for each FOV.

### Synthesis of HRO761

The WRN inhibitor HRO761 was synthesized following an established synthesis route for Compound 42, as described in WO2022/249060. (Bordas et al., 2022).

---

**Extended Data Fig. 1:**
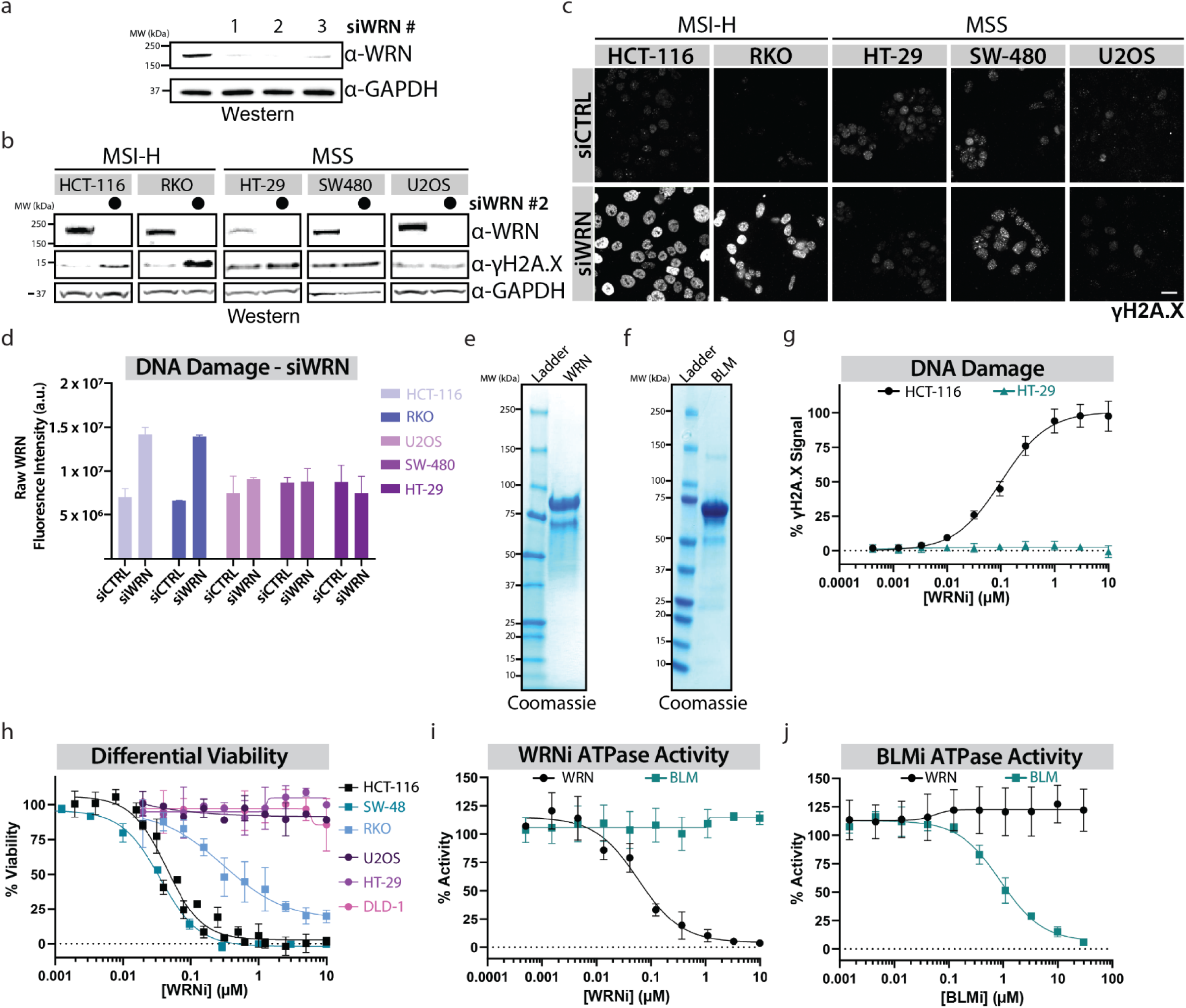
**a**. Whole cell lysates of HCT-116 after siRNA depletions of WRN for 24 h and probing with a WRN antibody show a robust loss of WRN protein. **b**. WRN depletion is synthetic lethal in MSI-H cells but not MSS cells. Whole cell lysates in the indicated cell lines after treatment with siWRN or siCTRL oligos for 48 h, and subsequently analyzed for DNA damage induction by Western blot. **c**. As in **b**, but cells were fixed in paraformaldehyde after siRNA treatments, and DNA damage was measured by measuring γH2A.X levels via immunofluorescence. Scale bar = 20 μm. **d**. Quantifications of **c.** Each graph represents the mean of *n* = 3 plates. **e**. Purification of WRN protein from SF9 insect cells. Coomassie gel staining shows a product of the expected protein molecular weight after purification. **f**. Protein purification of BLM protein from *E. coli*. Coomassie gel staining shows a product of the expected protein molecular weight after purification. **g**. DNA damage induction in MSI-H cells after WRN inhibition is dose-dependent. Dose response curves measuring DNA damage response via γH2A.X levels in HCT-116 cells or HT-29 cells after treatment with WRNi for 24 h. Graphs represent averages from *n* = 6 plates. **h**. Cell viability panel of MSI-H and MSS cells showing the differential viability effect of WRN inhibition towards MSI-H cells. Dose response curves measuring the viability of the indicated cell lines after WRNi treatment for 4 days. HCT-116, SW-48, and RKO are MSI-H cells; U2OS, HT-29, and DLD-1 are MSS cells. Graphs represent averages from *n =* 3 plates. All error bars represent standard deviation (s.d.). **i**. Dose response curves measuring the *in vitro* ATPase activity of WRN or BLM after WRNi treatment. Graphs represent averages from *n* = 6 plates. All curve fits were done by fitting a 4-parameter logarithmic regression curve. **j.** Purified BLM protein is active. Benchmarking of BLM protein by treatment with BLMi. Dose responses measuring ATPase and helicase inhibition by BLMi. All curve fits were done by fitting a 4-parameter logarithmic regression curve. All error bars represent standard deviation (s.d.). DMSO is dimethyl sulfoxide; WRNi is HRO761; BLMi is the BLM inhibitor Compound 2. MW is molecular weight.

**Extended Data Fig. 2:**
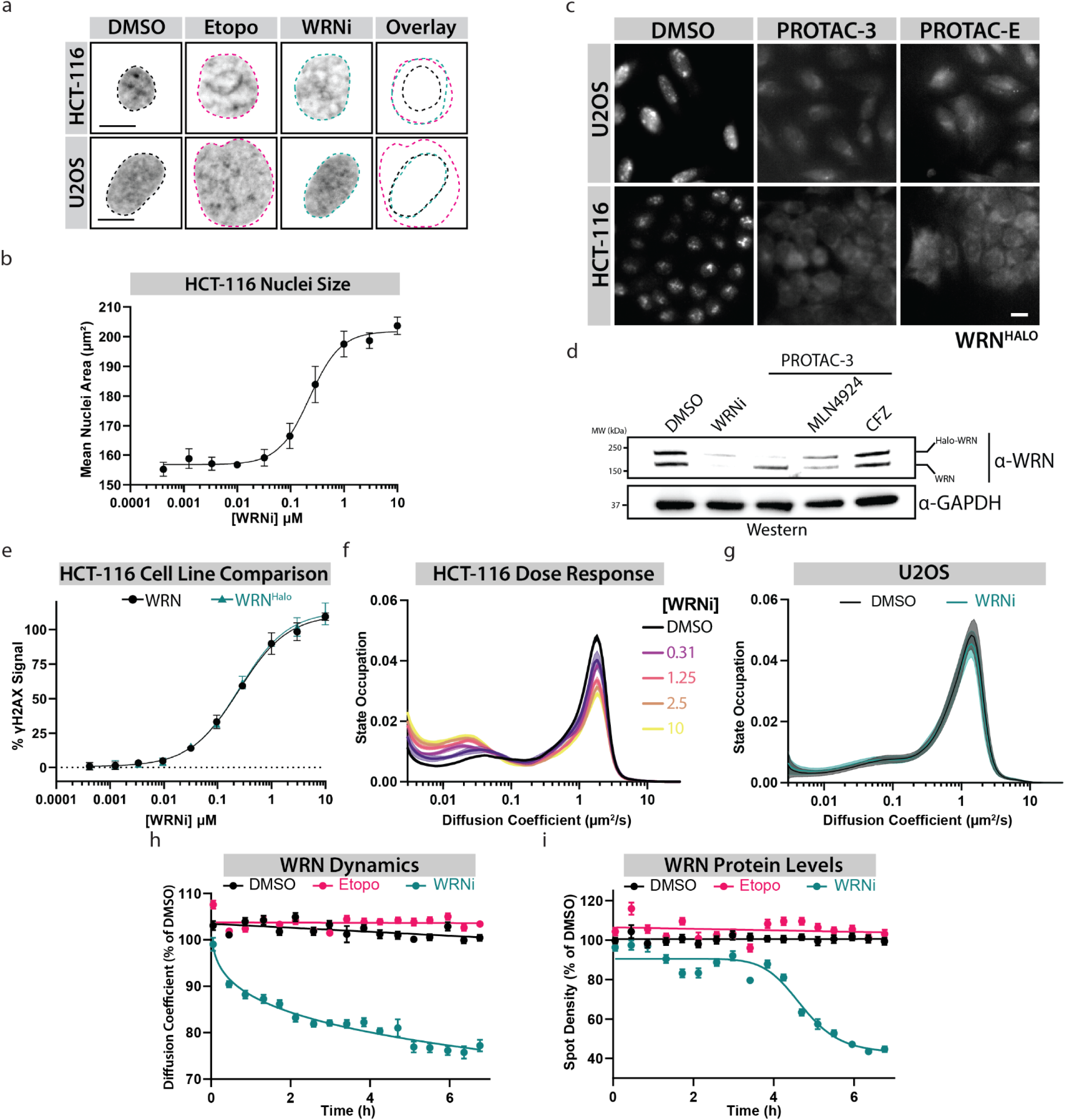
**a**. WRNi causes morphological changes to nuclei due to DNA damage accumulation in MSI-H cells. Images of Hoechst-stained nuclei of HCT-116 or U2OS cells treated in the presence or absence of 10 μM WRNi or etoposide for 24 h. Nuclei outlines are overlaid on top of each other, showing the large change in area after WRNi treatment in HCT-116 cells. This increase in nuclei size is observed in HCT-116 and U2OS cells after etoposide treatment. Scale bar = 10 μm. **b**. Nuclear morphology changes induced by DNA damage are dose-dependent. Dose response curve measuring nuclei area in HCT-116 cells after treatment with WRNi for 24 h. Error bars represent s.d.. **c**. Validation of endogenous WRN Halo tagging of HCT-116 and U2OS. WRN protein levels in HCT-116^WRN-Halo^ and U2OS^WRN-Halo^ visualized by staining with JF549 dye after treatment with 10 μM degraders of HaloTag, PROTAC-E or PROTAC-3, for 24 h. Scale bar = 10 μm. **d**. Treatment with Halo-PROTAC-3 leads to proteasomal dependent degradation of WRN^Halo^. Further validation of the WRN^Halo^ tag, showing Western blot analysis of HCT-116-WRN^Halo^ cells after treatment with 10 μM PROTAC-3 in the presence or absence of 2 nM CFZ or 5 nM MLN-4924 for 24 h. PROTAC-3 uses CUL2^VHL^ as a ligase, therefore CUL2 inhibition via MLN-4924 leads to a rescue of the WRN^Halo^ degradation phenotype. **e**. Wild type HCT-116 cells and HCT-116-WRN^Halo^ cells have identical responses to WRNi, suggesting WRN^Halo^ is functional. Dose response curves of HCT-116^WT^ and HCT-116-WRN^Halo^ measuring DNA damage induction. Error bars represent s.d.. **f.** WRNi leads to a dose-dependent increase of WRN molecules bound to chromatin. Distribution of diffusive states for WRN^Halo^ in increasing concentrations of WRNi. WRNi leads to a decrease of free-disusing WRN molecules with a concomitant increase in chromatin-bound WRN molecules. **g**. WRNi does not lead to chromatin trapping in MSS cells. SMT measurements show that the distribution of WRN^Halo^ diffusive states in U2OS^WRN-Halo^ cells remains unchanged in the presence or absence of 10 μM WRNi. Shaded area represent s.d.. **h** and **i**. WRN diffusion coefficient (**h**) and protein levels (**i**) remain unchanged upon DNA damage induction. Kinetic SMT of HCT- 116-WRN^Halo^ after treatment with 10 μM WRNi or Etopo over the indicated time points. Error bars represent s.e.m.. CFZ is carfilzomib; DMSO is dimethyl sulfoxide; WRNi is HRO761; Etopo is etoposide. MW is molecular weight.

**Extended Data Fig. 3:**
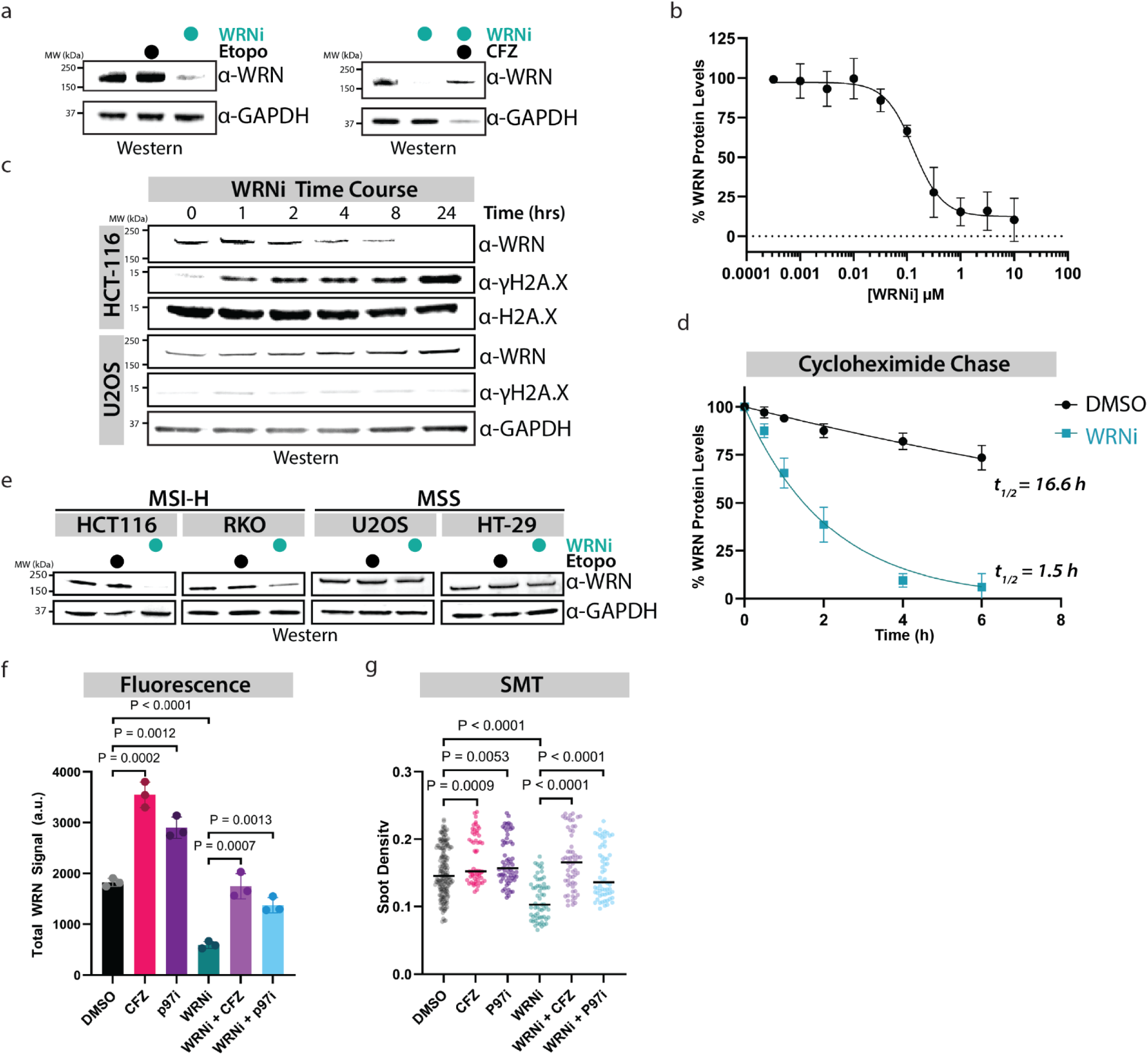
**a.** Degradation of WRN is induced by WRNi, but not by general DNA damage. Western blot analysis of HCT-116 cells treated with 10 μM etoposide or WRNi. This degradation is rescued by the addition of CFZ. **b**. Dose response curves measuring WRN protein levels by staining HCT-116^WRN-Halo^ cells with JF549 after treatment with WRNi for 24 h. Graphs represent averages from *n* = 3 plates, measuring 3 well per plate and 6 FOVs per well. The curve fit was done by fitting a 4-parameter logarithmic regression curve. **c**. Inhibition of WRN induces WRN degradation in a time-dependent manner in MSI-H cells. Steady state Western blot analysis of HCT-116 or U2OS cells treated with 10 μM WRNi over the indicated time points. d. Quantification of Fig. 3b. WRN inhibition leads to a decrease in the half-life of WRN protein. Cycloheximide (CHX) chase experiments in HCT-116 cells in the presence or absence of 10 μM WRNi show a dramatic decrease in the half-life of WRN protein upon its inhibition. Graphs represent *n =* 2 replicates. The curve fit was done by fitting a half-life decay regression curve. e. WRN degradation upon its inhibition is MSI-H dependent. The indicated MSI-H or MSS cell lines were treated as in a and analyzed by Western blot with the indicated antibodies. f. Quantifications of Fig. 3i. Bar graphs are the mean of n = 3 replicates, each point represents an the average of a well. Error bars represent s.d. g. SMT can be used to measure protein degradation. SMT was used to measure WRN molecules after inhibition of the p97/VCP-proteasome pathway, showing a rescue in protein degradation. Each point represents the average WRN spot density within all the nuclei in an FOV. n = 4 plates. Lines represent sample medians. DMSO is dimethyl sulfoxide; CFZ is carfilzomib; p97i is CB-5083; WRNi is HRO761. P-values were calculated using a two-tailed, unpaired Student’s t-test. MW is molecular weight.

**Extended Data Fig. 4:**
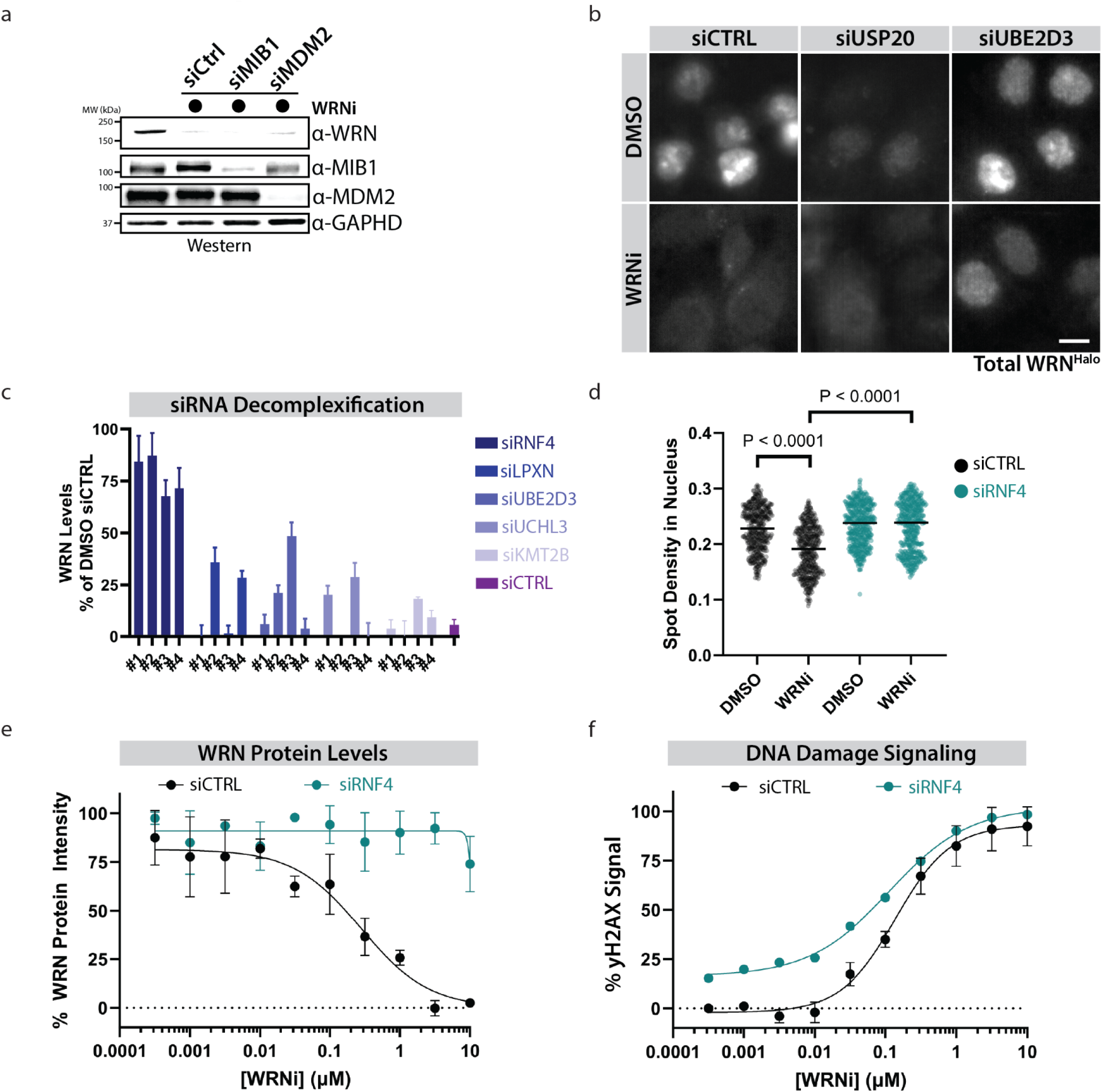
**a**. Previously reported E3 ligases of WRN are not responsible for the WRNi-dependent degradation phenotype. HCT-116 were treated with indicated siRNA oligos for 24 h, and subsequently treated with 10 μM WRNi for 16 h and analyzed by Western showing that depletion of the indicated E3 ligases does not rescue the WRN degradation phenotype. **b.** Identification of additional ubiquitin pathway regulators of WRN regulation. Representative images of HCT-116 cells after depletion of the indicated ubiquitin pathway regulators. Scale bar = 10 μm. **c**. Quantification of decomplexified siRNA screen hits from **Fig. 4a** after treatment of HCT-116^WRN-Halo^ with indicated siRNA oligos for 24 h and then subsequent treatment with 10 μM WRNi for 24 h. Bar graphs are the average quantified WRN protein levels for each indicated siRNA oligo of *n* = 3 plates. Error bars represent s.d.. **d**. SMT can be used to quantify protein degradation. Depletion of RNF4 rescues WRN protein levels after WRN inhibition. WRNi dot plots showing the WRN nuclear spot density from SMT experiments after co-treatment with siRNF4 and either DMSO or WRNi. Each point represents the average WRN spot density within all the nuclei in an FOV. n = 4 plates. Lines represent sample medians. **e**. RNF4 depletion rescues the WRN degradation phenotype. Dose response curves measuring WRN protein levels by imaging HCT-116^WRN-Halo^ treated with the indicated siRNAs for 24 h, and subsequent treatment with WRNi for 24 h. Graphs represent averages from *n* = 3 plates, measuring 3 wells per plate and 6 FOVs per well. Error bars represent s.d.. **f**. Depletion of RNF4 exacerbates DNA damage induced by WRNi. HCT-116 cells were depleted with indicated siRNAs for 24 h, and subsequently subjected to a dose response of WRNi for 16 h. DNA damage was assessed by measuring γH2A.X staining. Graphs represent averages from *n* = 3 plates, measuring 3 wells per plate and 6 FOVs per well. Error bars represent s.d.. All curve fits were done by fitting a 4-parameter logarithmic regression curve. DMSO is dimethyl sulfoxide; WRNi is HRO761. P-values were calculated using a two-tailed, unpaired Student’s t-test. MW is molecular weight.

**Extended Data Fig. 5:**
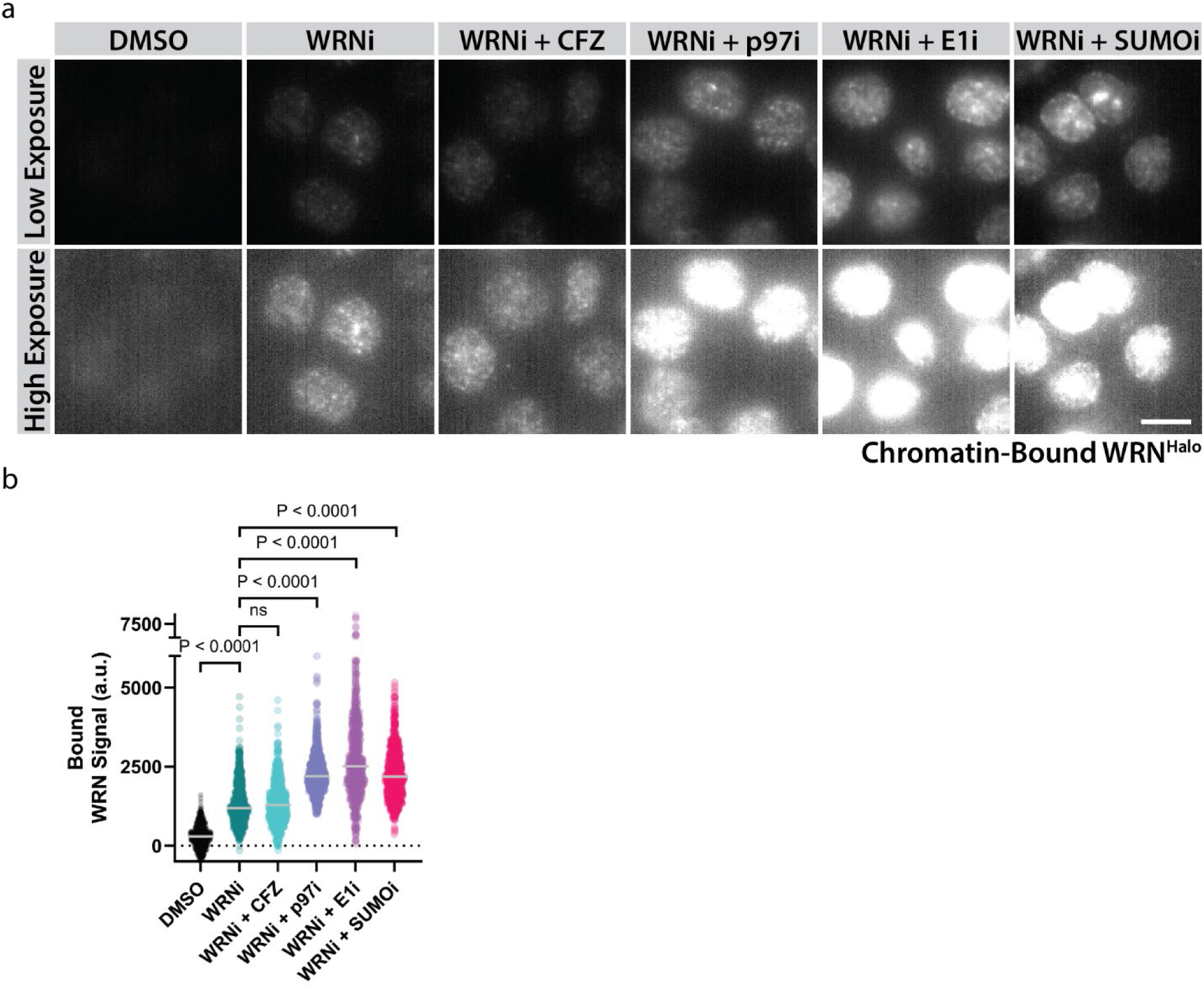
**a.** The SUMO-Ubiquitin-p97/VCP axis is required to remove trapped WRN from chromatin. Treatment of HCT-116-WRN^Halo^ cells with 10 μM WRNi in the presence or absence of 1 μM of CFZ, p97i, E1i, or SUMOi, followed by detergent extraction and imaging. **b.** Dot plot quantification of **a.** Each point represents an individual cell, measuring the nuclear intensity of WRN. Bars represent the means of the populations. DMSO is dimethyl sulfoxide, WRNi is HRO761; CFZ is carfilzomib; p97i is CB-5083; E1i is TAK-243; SUMOi is ML-792. P-values were calculated using a two-tailed, unpaired Student’s t-test. ns = not significant.

**Extended Data Fig. 6:**
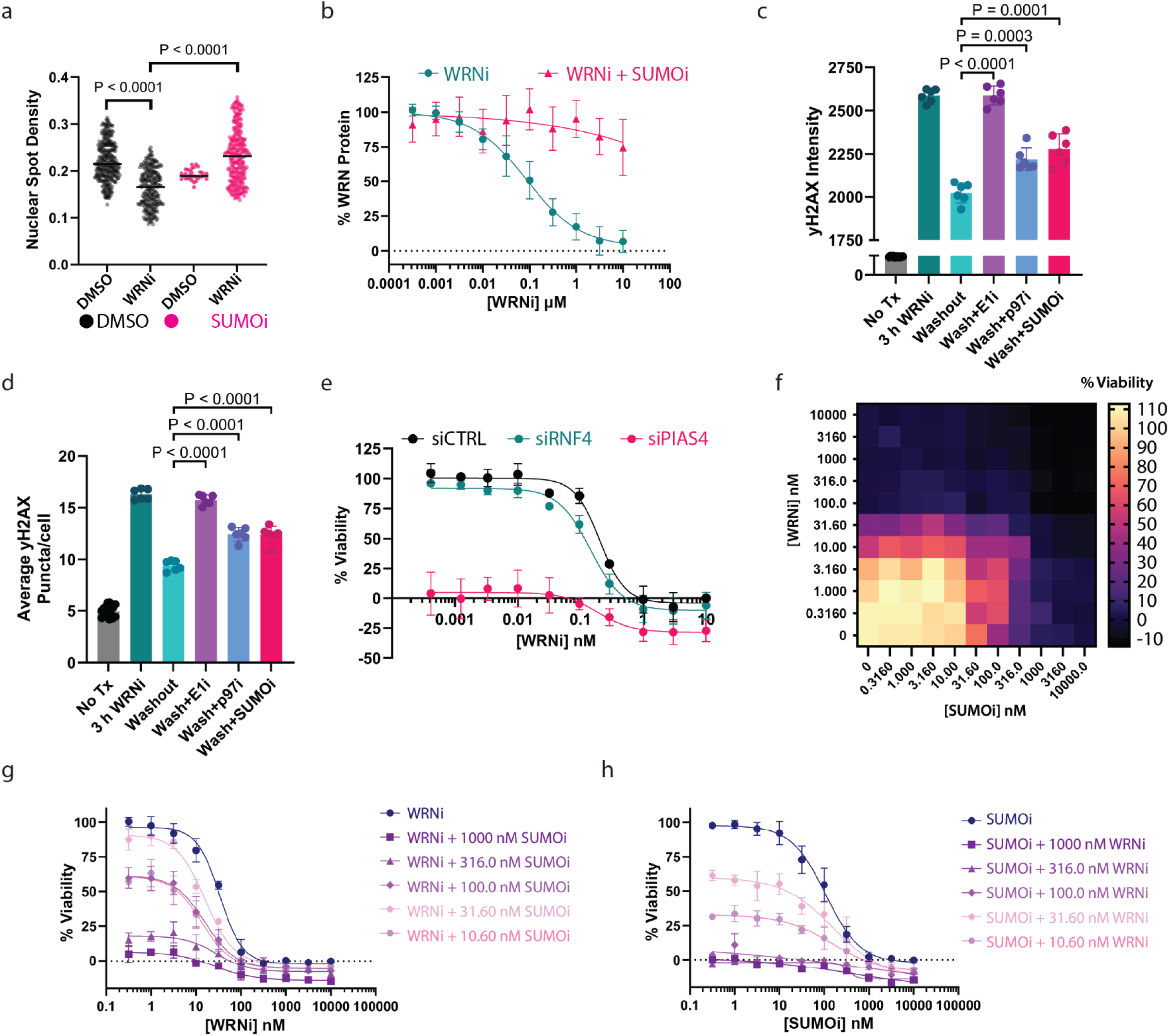
**a**. SMT can be used to elucidate molecular regulatory pathways, such as protein degradation. Inhibition of SUMOylation rescues WRN protein levels after WRN inhibition. WRNi Dot plots showing the WRN nuclear spot density from SMT experiments after co-treatment with SUMOi and either DMSO or WRNi. Each point represents the average spot density within all the nuclei in an FOV. n = 4 plates. Lines represent sample medians. **b**. Inhibition of SUMOylation prevents WRN degradation by WRNi. Dose response curves measuring WRN protein levels by in HCT-116^WRN-Halo^ treated with WRNi in the presence or absence of 1 μM SUMOi, and subsequent treatment with WRNi for 24 h. Graphs represent averages from *n* = 3 plates, measuring 3 well per plate and 6 FOVs per well. Error bars represent s.d. **c** and **d**. Quantification of washout experiment in **Fig. 5l**. **d** quantifies the average γH2A.X nuclear intensities of an FOV, and **c** quantifies the average number of γH2A.X puncta per nuclei in an FOV. Bar graphs in both **c** and **d** represent the average of *n =* 3 plates. Each data point represents the average of one well, containing 6 FOVs. Error bars represent s.d.. **e**. WRNi compound efficacy is independent of WRN degradation but shows potential sensitization to inhibition of the PIAS4-RNF4 axis. HCT-116 cells were treated with the indicated siRNAs for 24 h, followed by WRNi treatment at the indicated doses for 48 h. Cell viability was measured using a CTG2 kit. Graphs represent averages from *n* = 3 plates. Error bars represent s.d. **f**. Co-treatment of WRNi with SUMOi has potential synergy. Dose response matrix of both WRNi and SUMOi with indicated concentrations of compound. HCT-116 cells were treated with indicated dose combinations for 48 h. Viability was measured via CTG2. **g** and **h**. Quantifications of **f**, showing dose response plots for the indicated concentration combinations. Graphs represent averages from *n* = 3 plates. Error bars represent s.d.. All curve fits were done by fitting a 4-parameter logarithmic regression curve. DMSO is dimethyl sulfoxide; WRNi is HRO761; E1 is TAK-243; SUMOi is ML-792; p97i is CB-5083. P-values were calculated using a two-tailed, unpaired Student’s t-test.

