## Supplementary Figures for "WRN Inhibition Leads to its Chromatin-Associated Degradation Via the PIAS4-RNF4-p97/VCP Axis"

Supplementary Fig 1. Proton NMR of Cmpd 1

a

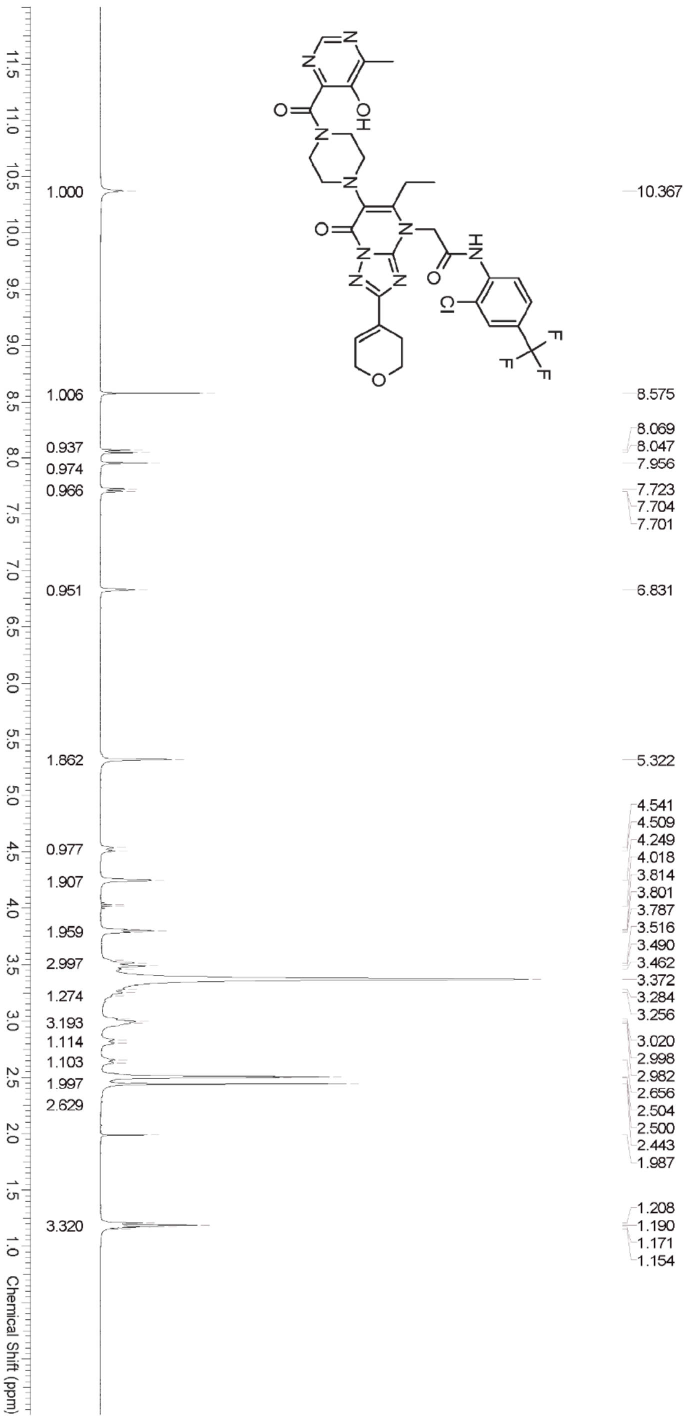


**Supplementary Fig. 1:**

$^{1}$H NMR spectrum was recorded on a Bruker ADVANCE spectrometer at 400 MHz. The chemical shifts are given in parts per million (ppm) on a delta (δ) scale. The solvent peak DMSO-$d_6$ = 2.50 ppm was used as a reference value for $^{1}$H NMR.

Supplementary Fig 3. LCMS characterization of Cmpd 1

a

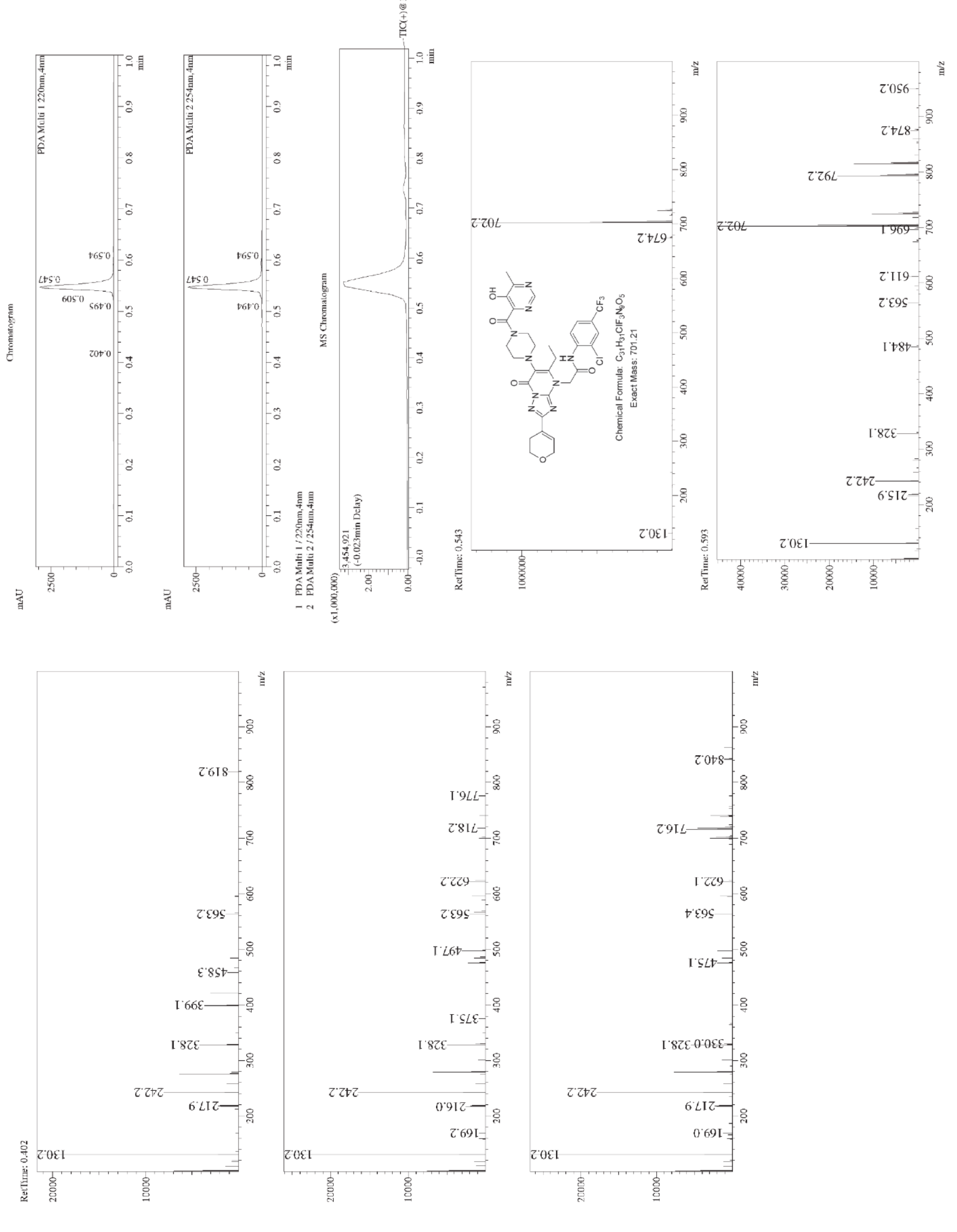


**Supplementary Fig. 2:**

Molecular weight of Compound 1 was confirmed by liquid chromatography mass spectrometry (ESI+) calcd for $C_{31}H_{31}ClF_3N_9O_5$ [M + H]+ 702.094, found 702.2.

Supplementary Fig 3. HPLC characterization of Cmpd 1

a

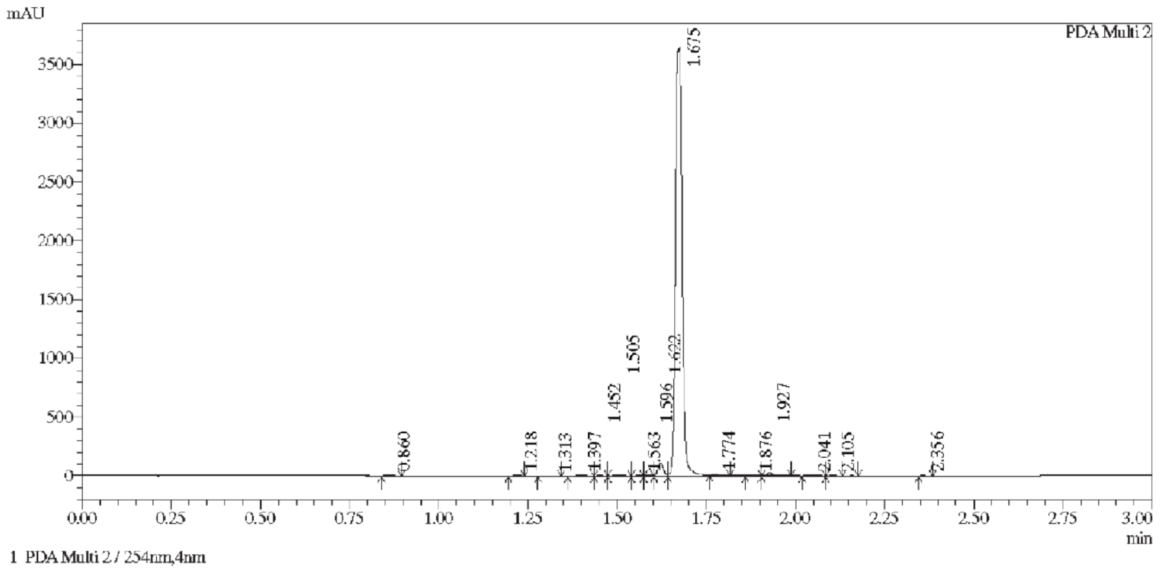


**Supplementary Fig. 3:**
Purity of Compound 1 was determined to be >95% by high performance liquid chromatography.
